# Cell size control emerges from the vein-dependent coordinated divisions of distinct cell groups in *Drosophila* wing

**DOI:** 10.64898/2025.12.26.696565

**Authors:** Kaoru Sugimura, Ryu Takayanagi, Toshinori Namba, Zeping Qu, Shuji Ishihara

## Abstract

During morphogenesis, cell divisions are precisely regulated in space and time. The biological objectives achieved by such regulation are not fully understood. Here, by applying a newly developed lineage-reconstruction pipeline to *Drosophila* pupal wing, we reveal that the wing is composed of distinct cell groups that differ in division number, timing, and spatial positioning relative to wing veins. We show that the frequencies of these lineages, together with their initial cell sizes and growth profiles, converge to achieve a highly conserved average cell size. Our data further suggest that distance from veins provides spatial information that biases where distinct lineages arise, and that loss of veins caused by perturbation of EGFR signaling suppresses a specific lineage and disrupts cell-size control. Finally, our results point to a multiscale organization of division patterns, in which vein-associated spatial information is integrated with local neighbor effects in a manner that would mitigate mechanical instability within the tissue. Together, these findings delineate a cell-size control mechanism based on coordinated divisions of distinct cell groups that supports robust morphogenesis and functional tissue design.

## Introduction

Cells proliferate to grow tissues, bend them through buckling, increase cell numbers, and perform other morphogenetic functions (Godard and Heisenberg, 2019; Hamant and Saunders 2020; Penzo-Méndez and Stanger, 2015; Vollmer et al., 2017; Irvine and Shraiman, 2017; Harmansa and Lecuit, 2021; Erlich and Harmansa, 2025; Tozluoǧlu and Mao, 2021). For these morphogenetic processes to proceed, diverse biochemical signaling pathways and mechanical cues trigger cell divisions (Duronio and Xiong, 2013; Gupta and Chaudhuri, 2022). These division-patterning mechanisms must operate in the context of dynamic and continuous tissue deformation while still encoding robust spatial information. In addition, they must function in ways that protect tissues from mechanical instabilities, such as unwanted tissue buckling (Tozluoǧlu and Mao, 2021). Despite their importance, the mechanisms underlying such robust control of cell divisions are still unclear. Moreover, because morphogenesis ultimately leads to the construction of a functional body, the biological objectives that cell-division regulation is optimized to achieve are of fundamental importance; however, they remain largely unexplored.

In multicellular tissues, tissue and cell size are primarily controlled by cell division and cell growth. Tissue size, cell division, and cell growth can be tightly coupled or largely independent, depending on genetic and environmental contexts (Potter and Xu, 2001; Penzo-Méndez and Stanger, 2015; Vollmer et al., 2017; Kuzmicz-Kowalska and Kicheva, 2021). At the level of individual cells, cell-autonomous mechanisms of cell-size control have been proposed, based on studies in unicellular organisms and *in vitro* cultured cells (Zatulovskiy and Skotheim, 2020; Lloyd, 2013; Ginzberg et al., 2015; Devany et al., 2023). In these systems, cells typically adjust the length of the G1 phase to grow toward a target size, or gradually converge on a characteristic size through repeated cell cycles via size-independent mass accumulation. The extent to which these mechanisms of cell-size control operate within animal tissues, however, is only beginning to be explored (Xie and Skotheim, 2020). In developing tissues, proliferation windows are often temporally restricted by systemic cues such as hormonal signals (Morrow and Mirth, 2024). Moreover, the timing of morphogenesis is coordinated across neighboring tissues, such that inter-tissue mechanical interactions can trigger or modulate morphogenetic processes (Collinet et al., 2015; Aigouy et al., 2010). How cell-size control through division and growth is adapted to such developmental constraints awaits further investigation.

*Drosophila* pupal wing serves as an excellent model system for studying tissue-scale coordination of cell division, owing to its reproducible development and access to whole-tissue imaging (Matamoro-Vidal et al., 2015; Diaz-de-la-Loza and Thompson, 2017; Fig. S1). During early pupal development, wing cells are arrested in the G2 phase through the action of the steroid hormone ecdysone (Guo et al., 2016; Morrow and Mirth, 2024). Cell divisions resume at ∼14–15 h after puparium formation (h APF), following the decline of the ecdysone pulse (Guo et al., 2016; Milán et al., 1996). Cells then undergo one or two rounds of division until ∼24 h APF, when a subsequent rise in ecdysone levels terminates proliferation, thereby restricting the division window to roughly 10 hours. Previous studies have measured several aspects of cell division in the pupal wing, including the orientation of divisions and the local tissue deformation produced by cumulative divisions (Garícia-Bellido et al., 1994; Milán et al., 1996; Aigouy et al., 2010; Etournay et al., 2015; Guirao et al., 2015; Trinidad et al., 2025). However, the mechanisms that determine where, when, and how many times cells divide during pupal development are yet to be fully elucidated. Furthermore, because the size of the wing blade is predetermined by the body size and remains nearly constant during the 10 h proliferation window, and because inhibiting cell division at this stage does not alter tissue size (Gokhale and Shingleton, 2015; Vollmer et al., 2017; Diaz-de-la-Loza et al., 2018; Aigouy et al., 2010; Weigmann et al., 1997), the functional significance of pupal-stage cell divisions remains elusive.

Here, we sought to elucidate how the number, timing, and positioning of cell divisions are regulated in the *Drosophila* pupal wing, and what these regulations are designed to achieve. By reconstructing individual cell lineages from whole-tissue time-lapse movies, we revealed that the pupal wing consists of distinct groups of cells that differ in their division number, division timing, and spatial distribution. The temporal profiles of these distinct division types are highly reproducible across samples, such that the total number of divisions is precisely tuned to maintain a target average cell size. Furthermore, we demonstrate that the spatio-temporal coordination of cell division and the precise control of average cell size depend on vein-associated epidermal growth factor receptor (EGFR) signaling.

## Results

### Pipeline to reconstruct cell-division lineages from whole-wing movie data

To investigate the spatiotemporal coordination of cell divisions in the pupal wing, we developed a pipeline that allows tracking of cell divisions during wing development (Fig. 1A, S1A–C; Materials and Methods). Using a *DE-cadherin (DE-cad)-GFP knock-in* line, in which the endogenous DE-cad protein is fused to GFP and expressed under its native promoter (Huang et al., 2009), we performed whole-wing time-lapse imaging from 15.5 h APF to 32 h APF. Individual cell division events occurring within the region of interest (ROI) were identified through cell tracking. By integrating cell tracking and division data, we reconstructed the lineage trees of individual cells, hereafter referred to as division trees, with associated temporal and positional information of each division event (Fig. 1B). Combining data from three wild-type wings, we observed over 10,000 cell divisions in the wing-blade ROIs.

**Figure 1.** Distinct groups of cells differing in division number and timing. (A) Flowchart of the data analysis. Whole-wing images were acquired at 5-min intervals starting from 15.5 h APF. Images were segmented and skeletonized using custom-made macros and plugins in ImageJ/Fiji (Ishihara and Sugimura, 2012; Guirao et al., 2015) or **EpySeg** (Aigouy et al., 2020). From the skeletonized images, cell geometry data, including vertex positions and connectivity, were extracted using the ImageJ/Fiji plugin **GetVertex**. This cell geometry data was directly used as input for Bayesian force inference (BFI; Ishihara and Sugimura, 2012). Cell tracking identified correspondences of cells between successive frames and detected cell division events. The lineage trees of individual cells are reconstructed by integrating cell tracking and division data. See Materials and Methods for details. (B) Division events were classified based on the structure of the division tree. Divtype 1 represents a division that produces two daughter cells from a single root cell (green). Divtype 2 includes events that result in the production of granddaughter cells (orange). Divtype 2 is further subdivided into: Divtype 2a, where one daughter and two granddaughter cells are produced (magenta), and Divtype 2b, where four granddaughter cells are produced (orange). (C) Frequencies of each division type are plotted over developmental time (green: divtype 1; orange: first division in divtype 2; light orange: second division in divtype 2). The frequency of each division type at a given time point was calculated as the number of divisions of that type occurring at that frame, divided by the total number of divtype 1 and divtype 2 divisions observed over the entire analysis period. (D) Snapshots from Movie 1. Cells belonging to lineages that existed at the initial time point are color-coded according to their division type. Green, divtype 1; orange, divtype 2; gray, other cells within the ROI, white: cells outside the ROI. Colors fade with successive divisions. Developmental time is indicated in the top-right corner. Data are plotted as mean ± SD (C). n = 3 wild-type wings (C).

### Wings cells are categorized into groups differing in division number and timing

The newly developed pipeline allows classification of cell lineages based on the structure of their division trees. Division type 1 (hereafter referred to as divtype 1) corresponds to lineages that undergo a single round of division, whereas division type 2 (divtype 2) undergoes two rounds (Fig. 1B). Our analysis uncovered that these two division types differ not only in the number of divisions but also in their temporal dynamics: the peak of divtype 1 divisions occurs around 18–19 h APF, between the two peaks of divtype 2 divisions (Fig. 1C). Furthermore, cells belonging to each division type tend to occupy distinct regions of the wing (Fig. 1D and Movie 1). The temporal profiles of these division types were highly reproducible across samples without the need for time alignment (Fig. S2A–C), indicating that division types are tightly regulated during the wing development.

### Temporal profiles of average cell size are highly conserved, reflecting those of division types

We next examined the consequences of the precisely regulated division types. By definition, different division types produce different numbers of progeny (Fig. 1B): divtype 1 lineages generate two daughter cells from a single “root” cell, whereas divtype 2 lineages are subdivided into divtype 2a, which yields one daughter and two granddaughter cells, and divtype 2b, which yields four granddaughter cells. Because the total tissue size changes only minimally (the size of the wing-blade ROI at 32 h APF is 1.04 ± 0.04 times that at 15.5 h APF in n = 3 wild-type wings), the average cell size, calculated as tissue size divided by cell number, can be strongly influenced by the frequencies of division types. Consistent with this, our measurements of cell size, defined as the apical cell area at the adherens junction plane, showed that the temporal profiles of average cell size are highly conserved across samples (Fig. 2A; Fig. S3). This conservation of average cell size is reflected in the tight association between tissue size and total cell number (black circles in Fig. 2E, F). Furthermore, the total number of progeny cells generated by division types 1 and 2 up to 24 h APF is strongly correlated with tissue size at this time point (black circles in Fig. 2G), suggesting that apoptosis or the fraction of non-dividing cells contribute negligibly. These findings suggest that the total number of divisions is tuned to maintain a target average cell size by regulating the frequencies of division types.

**Figure 2.** Conserved profiles of cell size and their dependence on division types. (A) Average cell area of all cells within the wing-blade ROI at each time point. (B) Time profiles of average area for cells undergoing divtype 1 (green) and divtype 2 (orange) divisions. (C) Time profiles of average area for cells undergoing divtype 1 (green), divtype 2a (magenta), and divtype 2b (orange) divisions. (D) Summary of results from Figs. S4 and S5, highlighting how temporal profiles of average cell size depend on division types and division tree layers. Numbers indicate the aligned time points for analysis (1, root cells at 60 min before the first division; 2, daughter cells at 60 min after the first division; 3, daughter cells at 60 min before the second division; 4, granddaughter cells at 60 min after the second division; 5, granddaughter cells at 240 min after the second division). In Fig. S5C, D, data from 20–5 min before division were excluded (Supplementary Information). Consequently, the area increase between time points 2 and 3 in divtype 2a and divtype 2b cells reflects interphase cell growth rather than cell expansion associated with mitotic cell rounding. (E, F) Tissue size plotted against the cell number at 15.5 h APF (E) and 24 h APF (F) (black circle: wild type, magenta diamond; veinless, gray square: *ds* RNAi). (G) Tissue size at 24 h APF plotted against the number of total progenies produced by divtype 1 and divtype 2 divisions up to 24 h APF (black circle: wild type, magenta diamond; veinless, gray square: *ds* RNAi). Data are plotted as mean ± SD (A–C). n = 3 wild-type wings (A–C, E–G). n = 3 *ds* RNAi wings (E–G). n = 2 veinless wings (E–G).

In addition to the frequencies of division types, the average cell size is also shaped by the initial size and the size changes that occur following division in each division type. For divtype 1 cells, the initial size was approximately 30 µm², remaining in a plateau phase until ∼17 h APF before undergoing a monotonic decrease between 17–20 h APF, which coincides with their division window (green curve in Fig. 2B). Divtype 2 cells showed two phases of monotonic decrease separated by a plateau, again consistent with the timing of their divisions (orange curve in Fig. 2B). When divtype 2a and divtype 2b cells were analyzed separately, the plateau and second decrease phases were less distinct in divtype 2a cells, while the cell-growth phase was observed between two decreasing phases in divtype 2b cells (magenta and orange curves in Fig. 2C). To directly monitor changes in cell size before and after division, we aligned individual cell tracks in time relative to their division events. This analysis revealed that each division type differs in the initial size of its root cell and exhibits distinct cell growth dynamics (summarized in Fig. 2D; Fig. S4, S5; Supplementary Information). The root cells of divtype 2b, which undergo two rounds of division, were initially only ∼20% larger than those of divtype 1, which undergo a single division, but they gradually enlarged during interphase before the second division and continued to grow post-mitotically. Divtype 2a cells initially resembled divtype 1 cells in size, but cells that underwent a second division became progressively larger after the first division, ultimately exhibiting size dynamics similar to divtype 2b cells. Consequently, final cell sizes converged to similar values by 24 h APF, when cell divisions cease. In summary, our data demonstrate that precise control of average cell size emerges from the coordination of distinct cell groups through both their frequencies and temporal size profiles.

### Different division-type cells are positioned differentially with respect to wing veins

Having established how division types cooperate to control average cell size, we next examined how distinct division-type lineages are specified. In our search for factors coordinating division types, we found that the spatial distributions of divtype 1 and divtype 2 cells vary between samples, but nevertheless exhibit a consistent pattern (Fig. 1D, S2D–F, S7). Because the PD alignment of divtype 1 and 2 cells is reminiscent of the longitudinal veins L3 and L4, we simultaneously tracked cell divisions and putative vein cells (see Fig. S1A and Materials and Methods for vein nomenclature and retrospective labeling of putative vein cells). This analysis revealed that divisions tend to initiate in proximity to veins and that divtype 1 cells are enriched along longitudinal veins, whereas divtype 2 cells are located in the intervein regions adjacent to longitudinal veins (Fig. 3A–C, Movie 2). At 15.5 h APF, the fraction of cells belonging to divtype 2 lineage was 0.045 ± 0.004 among putative vein cells, compared with 0.24 ± 0.02 among putative intervein cells (n = 3 wild-type wings). To quantify these patterns, we measured the distance from each mother cell to the nearest vein cell. Fig. 3D shows that the shortest distance to veins increased over time for mother cells, albeit with large variance (blue line in Fig. 3D). As a control, the average distance for all cells remained nearly constant (gray line in Fig. 3D). Both divtype 1 and divtype 2 divisions followed this increasing trend, but divtype 1 divisions consistently occurred closer to vein cells than divtype 2 divisions at earlier time points (Fig. 3E). These observations indicate that different division-type cells are positioned differentially with respect to wing veins, suggesting a possibility that wing veins may contribute to regulating division types.

**Figure 3.** Spatial relationship between division-type groups and vein positions. (A) Snapshots from Movie 2, showing cells generated by divisions occurring within the ROI (yellow). Putative vein cells are shown in blue and light blue. Developmental time is indicated in the top-right corner. The arrows indicate an infrequent division around the PCV. (B, C) Cells are labeled with dots, with colors indicating their division type using the same color scheme as in Fig. 1D (B: 16 h APF; C: 24 h APF). Dot colors fade with successive divisions. Putative vein cells are overlaid in blue or light blue. The region outlined by a square is shown at higher magnification on the right. In (B), asterisks mark pairs of daughter cells, and the arrow indicates that divtype 2 divisions are infrequent in intervein regions abutting the PCV, in contrast to those abutting L4 or L5. (D, E) The distance from each cell to the nearest putative vein cell is plotted over time. Gray lines: all cells; blue lines: mother cells (defined as cells 5 min before division). In (E), mother cells are further separated by divtype: green lines for divtype 1 and orange lines for divtype 2. Data are plotted as mean ± SD (D, E). n = 3 wild-type wings (D, E).

### Discoordination of division timing, division type, and cell size in veinless wings

To directly test the relevance of veins in specifying distinct division lineages, we analyzed the veinless mutant, in which wing cells fail to differentiate into vein cells due to downregulation of EGFR signaling (García-Bellido et al., 1994; Blair, 2007; Fig. S1D). In veinless wings, the fraction of divtype 2 divisions decreased dramatically, with divtype 2b divisions being primarily affected, and the clear separation between divtype 1 and divtype 2 division peaks observed in wild type was lost (Fig. 4A, B). This decrease in divtype 2 divisions is consistent with the previously reported reduction in BrdU incorporation at 13–16 h APF, although that study did not discriminate among division types (García-Bellido et al., 1994). The spatial arrangement of different division-type cells was also disrupted and no longer apparent (Fig. 4C; Movie 3). Furthermore, various aspects of cell size control were altered upon vein loss. Veinless wings retained, to some extent, division-type-specific initial cell sizes and their temporal size profiles (Fig. 4F, G). However, these quantitative relationships were significantly shifted, resulting in 30% increase in average cell size compared with wild-type intervein cells at 24 h APF (Fig. 4E–H). Consequently, changes in the fractions of division types and their temporal size profiles led to clear deviations in both the tissue size-cell number and tissue size-progeny number relationships (magenta diamonds in Fig. 2F, G). Interestingly, such deviations were not observed at 15.5 h APF (Fig. 2E). Thus, although wing size and the eventual division-type composition have already diverged from wild type by this stage, significant disruption of tissue-level control of average cell size becomes evident only later, after the period of active proliferation. Together, these results indicate that veinless mutations, known to perturb EGFR signaling and vein fate, disrupt the coordination of division types and cell-size regulation during pupal wing development, and that proper coordination among distinct division types is a key mechanism for tuning average cell size.

**Figure 4.** Discoordination of division timing, division type, and cell size in veinless wings. (A) Frequencies of each division type plotted over developmental time (green: divtype 1; orange: first division in divtype 2; light orange: second division in divtype 2). (B) Table showing the number of progeny produced by divtype 1, divtype 2, divtype 2a, and divtype 2b divisions up to 24 h APF, normalized by the total number of cells within the wing-blade ROI at 15.5 h APF, for the indicated genotypes. (C) Snapshots from Movie 3 showing cells color-coded by divtype, as presented in Movie 1 and Fig. 1D. (D) Snapshots from Movie 6, showing cells generated by divisions within the ROI, as in Movie 2 and Figure 3A, but without the overlay of vein cells. (E) Average cell area of all cells within the wing-blade ROI at each time point. (F) Time profiles of average area for cells undergoing divtype 1 (green) and divtype 2 (orange) divisions. (G) Time profiles of average area for cells undergoing divtype 1 (green), divtype 2a (magenta), and divtype 2b (orange) divisions. (H) Table of average cell size for all cells, putative intervein cells, and putative vein cells at 24 h APF for the indicated genotypes. No vein cells are present in veinless mutant wings. Data are shown as mean ± SD (A, B, E–H). n = 3 wild-type wings (B, H). n = 2 veinless wings (A, B, E–H).

The pupal wing develops planar cell polarity (PCP) along the proximal-distal (PD) axis, specified by counter-gradients of Dachsous (Ds) and Four-jointed (Butler and Wallingford, 2017; Strutt and Strutt, 2021). In addition, wing cells elevate tension at PD-oriented junctions to resist extrinsic pulling forces from the hinge (Aigouy et al., 2010; Sugimura and Ishihara, 2013). These biochemical and mechanical fields within the tissue could encode spatial information that coordinates cell-division patterns, such as division orientation. However, we found that although *ds* RNAi produced a more rounded wing with reduced PD elongation (Fig. 5C, D, S1E; Baena-López et al., 2005), division timing, division-type composition, and cell-size regulation were not significantly altered (gray squares in Fig. 2E–G; compare Fig. 5A with Fig. 1C, Fig. 5E–G with Fig. 2A–C, and Fig. 5I, J with Fig. 3D, E; Fig. 5B, H; Movies 4, 5). In addition, inferred junction tension showed no strong correlation with division types, and average tensions for junction-angle classes were indistinguishable among division types, both at 15.5 h APF and 20 h APF (Fig. S8). Taken together, while our *ds* RNAi and force inference analysis do not exclude contributions from PCP or mechanical forces, the results are overall consistent with a role for vein-associated EGFR signaling in coordinating distinct division types with their characteristic timing and numbers, thereby ensuring precise control of average cell size.

**Figure 5.** Division timing, division type, and cell size are not significantly altered in *ds* RNAi wings. (A) Frequencies of each division type are plotted over developmental time (green: divtype 1; orange: first division in divtype 2; light orange: second division in divtype 2). (B) Snapshots from Movie 4 showing cells color-coded by divtype, as presented in Movie 1 and Fig. 1D. Developmental time is indicated at the top-right corner. (C) Snapshots from Movie 5, where cells generated by divisions and putative vein cells are labeled, as presented in Movie 2 and Fig. 3A. (D) Average cell area of all cells within the wing-blade ROI at each time point. (E) Time profiles of average area for cells undergoing divtype 1 (green) and divtype 2 (orange) divisions. (F) Time profiles of average area for cells undergoing divtype 1 (green), divtype 2a (magenta), and divtype 2b (orange) divisions. (G, H) The distance from each cell to the nearest putative vein cell is plotted over time. Gray lines: all cells; blue lines: mother cells (defined as cells 5 min before division). In (H), mother cells are further separated by divtype: green lines for divtype 1 and orange liens for divtype 2. Data are plotted as mean ± SD (A, B, E–J). n = 3 wild-type wings (B, H). n = 3 *ds* RNAi wings (A, B, E–J).

### Multiscale organization of division patterns that integrate vein proximity and local neighbor effects

So far, we have focused on the regulation and biological consequences of the total number of cell divisions. Our division-tracking analysis also revealed that the spatial distribution of cell divisions follows a structured pattern during pupal wing development. Division events propagated from proximal to distal regions of the wing (Fig. 1D, 3A; Movies 1, 2). This spatial progression of divisions roughly followed the distance from vein cells: longitudinal veins, especially L2 and L5, are not perfectly parallel to the proximal-distal axis; instead, they converge in the proximal region, reducing the distance to veins there (Fig. 6A, B). The vein-distance map is largely conserved during the proliferation period (Fig. 6C), despite ongoing tissue deformation. The spatial positioning of veins, together with the vein distance-dependent division timing (Fig. 3D, E), may account for the observed global wave of cell-division events. A similar initiation of divisions was also observed near the wing margins, although these margin-associated divisions contributed less to the overall propagation pattern (see Supplementary Information for discussion). Cell divisions were infrequent around the posterior crossvein (PCV) (arrows in Fig. 3A). Unlike the other longitudinal veins and the anterior crossvein, the PCV is specified later, at around 20–24 h APF, via Decapentaplegic (Dpp) transport from neighboring longitudinal veins (Blair, 2007; Matsuda and Shimmi, 2012). In line with this, discontinuity of the L4 vein upon overexpression of Argos, a negative regulator of EGFR signaling (Blair, 2007), results in reduced cell division in the vicinity of L4 (Fig. S9). Moreover, the global pattern of cell division is disrupted in the veinless wing: two directed waves of cell division emerge from both the proximal and distal regions, and the onset of cell divisions in the central region is delayed until around 19–20 h APF (Fig. 4C, D; Movies 3, 6). Our observations are therefore consistent with the idea that acquisition of vein identity is linked to the global progression of cell divisions.

**Figure 6.** Multiscale organization of division patterns that integrate vein proximity and local neighbor effects. (A, B) Vein-distance maps at 15.5 h (A) and 24 h APF (B). The color bar indicates the distance to the nearest putative vein cell. (C) The distance from each cell to the nearest putative vein cell is plotted over time. Among tracks that persist from 15.5 to 24 h APF, those corresponding to vein cells and those with an initial distance ≥ 90 µm were excluded. When a cell division occurred, only one daughter cell was followed. The remaining tracks were grouped into three bins based on their initial distance to veins, and 10 tracks were randomly selected from each bin. Different colors indicate different bins. (D, E) Local progression of cell division events. (D) Cells generated by division are shown in yellow, and asterisks indicate newly divided cells in the current frame. Developmental time is indicated at the top-right corner. (E) Cells generated by divisions are color-coded based on their division orientation (see color bar at the bottom). (F, G) Same as (D) and (E), but from a different wild-type sample. (H, I) At each time point (frame *t*), cells were classified based on whether at least one of their neighbors had divided. We then examined whether cells in each group divided in the subsequent frame (*t* + 5 min). The division probability plotted at time *t* + 5 min was defined as the number of cells that divided between t and *t* + 5 min divided by the number of cells in the group at time *t*. Yellow lines represent cells with at least one divided neighbor; black lines represent cells with no divided neighbors. In (I), cells were further subdivided based on their distance from the nearest putative vein cell. Lighter-colored lines correspond to cells located 20 µm or more from the nearest putative vein cell. (J, K) A local cluster is defined as a group of six or more neighboring cells that have divided by a given time frame (birth time). For each cluster, the birth time and the minimum distance from the cluster centroid to the nearest vein cell were measured. Cells belonging to identified clusters were excluded from further analyses from the next frame onward. (J) Scatter plot showing the relationship between cluster birth time and the minimum distance from the cluster centroid to the nearest vein cell for divisions occurring at or before 18 h APF. Different colors indicate data from individual wild-type samples, each containing multiple clusters. (K) Histogram of the minimum distances between cluster centroids and the nearest vein cells, generated from the data shown in (J). Data are plotted as mean ± SD (H, I). n = 3 wild-type wings (H–K).

Beyond this global pattern, cell division events also propagate at the local scale to neighboring cells: we found that cells adjacent to recently divided cells were more likely to divide next, forming local “chains” of divisions (Fig. 6D, F; Fig. S10A, C). These division chains offer a live view of a process that could give rise to the clusters of *string* (*cdc25*)-expressing cells reported by Milán et al. (1996). Unlike the global progression of cell division, the orientation of local division chains is not aligned, and a single cluster of divided cells can expand in multiple directions, partly reflecting variation in division orientation (Fig. 6E, G; Fig. S10B, D). To quantify the neighbor effect in triggering division, we classified cells based on whether at least one neighbor had divided and then assessed whether those cells divided in the following frame (*i.e.*, 5 min later). As clearly shown in Fig. 6H, cells with divided neighbors showed a markedly higher division probability.

Strikingly, the neighbor effect exerted a stronger influence than vein distance: cells located more than 20 µm from veins (approximately the mean distance to veins for all cells; see Fig. 3D) but with divided neighbors had a higher division probability than cells close to veins but lacking divided neighbors (light yellow versus black lines in Fig. 6I). Moreover, a neighbor effect, albeit weaker, was also observed in the veinless wing (Fig. S11). Nevertheless, in wild-type wings, most clusters of divided cells were found near veins at early time points (Fig. 6J, K and their figure legends). Thus, veins bias the locations at which local clusters are born, rather than simply promoting individual divisions in a distance-dependent manner. Together, the global pattern of cell division emerges from the summation of undirected local chains of divisions triggered by neighbor effects and spatially biased by veins.

### Loose coupling between vein distance and division progression may preserve mechanical stability during proliferation

The cell-division patterns that combine vein-dependent seeding with neighbor-dependent spreading result in a loose coupling between division timing/position and vein distance. To investigate the biological significance of this loose coupling, we performed numerical simulations of cell-division progression using a Cell Vertex Model (Fletcher et al., 2014). Specifically, we compared three scenarios (Fig. S12A–C; Supplementary Information): (A) stochastic propagation of cell divisions, in which the first division events are initiated only in cells adjacent to veins and subsequently spread through the neighbor effect; (B) stochastic propagation of cell divisions, in which division events are initiated by vein-distance-dependent spontaneous seeding and spread through the neighbor effect (motivated by the experimental observations in Fig. 6H–K); and (C) deterministic progression of cell divisions strictly determined by vein distance, without neighbor-dependent spreading.

We found that, in scenario C, the average cell area normalized to its target value decreased as domains of divided cells expanded (Fig. S12D), suggesting increased cellular compression caused by mitotic cells with enlarged areas. Notably, once the number of mitotic cells exceeded a certain level, mean cell area dropped sharply in scenario C, whereas it remained relatively stable in the other scenarios at comparable numbers of mitotic cells (Fig. S12E). Finally, scenario B expanded division domains more efficiently than scenario A: the time required for division progression from 10% to 90% of cells undergoing division was shorter in scenario B (Fig. S12F). This difference arises because, in scenario A, divisions initiated only in vein-adjacent cells tend to propagate primarily in directions away from the vein. In contrast, in scenario B, spontaneous initiation of divisions in cells distant from veins enables multidirectional expansion of division domains. Collectively, these results suggest that the multiscale organization of division patterns revealed by this study may help mitigate mechanical instability within the tissue while maintaining efficient propagation of cell divisions.

## Discussion

By following over 10,000 cell divisions during *Drosophila* pupal wing development, this study identifies groups of cells that differ in division number, timing, and spatial distribution. Our data further demonstrate that the frequencies of these groups, together with their initial cell sizes and growth profiles, collectively achieve a highly conserved average cell size under a predetermined and nearly constant tissue size. Unlike established mechanisms of cell-size control, which rely on repeated cell cycles, extensive cell growth, or feedback from tissue size, this mechanism operates within a hormonally regulated, limited window of proliferation and associated interphase cell growth and instead exploits distinct division-type lineages. These findings reveal a novel strategy for cell-size control adapted to the developmental constraints of the system.

What is the advantage of having multiple division types rather than a single one? In the *Drosophila* wing, final tissue size is set at very early pupal stages in proportion to body size (Gokhale and Shingleton, 2015; Vollmer et al., 2017; Diaz-de-la-Loza et al., 2018). Accordingly, we consider tissue size to be conserved in the following discussion. If only divtype 2b cells were present, wing cells need to undergo two rounds of division within the 10-hour proliferation window. Given that the second cell cycle lasts ∼5 hours, as inferred from the interval between the first and second peaks of divtype 2 divisions, these two rounds of divisions would proceed nearly synchronously. Such synchronized divisions generate a surge of adhesion remodeling and mechanical stress, which epithelial tissues may not tolerate and could lead to tissue rupture (Pinheiro and Bellaïche, 2018; Godard and Heisenberg, 2019). Conversely, if only divtype 1 cells were present, maintaining the correct average cell size after the completion of the proliferation phase would require increasing the initial cell number (equivalently, the terminal cell number in the larval wing disc) by ∼25%. This would likely impose stronger compressive stresses on the larval wing pouch (Harmansa and Lecuit, 2021). Thus, the coexistence of distinct division types may help protect wing tissues from structural and mechanical instability.

How different division-type lineages are specified is yet to be fully elucidated. Our data indicate that in the veinless mutant wing, divtype 2 is selectively suppressed. The veinless mutant carries mutations in *vein*, which encodes an EGF ligand, and *rhomboid*, which is required for secretion of the active form of another EGF ligand, Spitz (De Celis, 1998; Shilo, 2005). Intricate interactions among the EGFR, Dpp, and Notch signaling pathways specify and refine the vein and intervein domains (Blair, 2007; Herszterg et al., 2025), which could influence the emergence of divtype 2 cells. However, although division types are preferentially positioned relative to veins, they do not map one-to-one onto cell fates. For example, while most vein cells undergo divtype 1 divisions, many divtype 1 cells are also distributed broadly throughout intervein regions. Moreover, the spatial organization of division types varies more across samples than the differentiation pattern itself, even when compared with pre-refined provein regions (Herszterg et al., 2025). The origin of this discrepancy and variability remains unknown, and it is not even clear to what extent division-type lineages correspond to EGFR/Dpp/Notch-regulated cell fates.

Regardless of the degree to which division types correspond to cell-fate specification, the mechanisms that determine the timing and number of divisions, as well as the size profiles within each division type, are not well defined. EGFR/ERK signaling promotes cell-cycle progression through transcriptional regulation and also influences cell growth (Lavoie et al., 2020). Cell-size-sensing mechanisms, such as Hippo signaling (Aragona et al., 2013; López-Gay et al., 2020; Zhong et al., 2024; Bosveld et al., 2026), can tune cell growth and cell-cycle progression and may bias larger cells toward undergoing additional (second) rounds of division. Mechanical properties may differ between vein and intervein cells, and spatially patterned mechanical environments can modulate cell proliferation (Martino et al., 2018). Vein cells are smaller than intervein cells, and their reduced size can stretch adjacent intervein cells; such stretch-induced enlargement may increase proliferative potential (Gudipaty et al., 2017; Gupta and Chaudhuri, 2022) and favor divtype 2 divisions in those adjacent intervein cells. Space- and time-resolved analyses of forces and stresses, beyond the averaged junction-tension measurements performed here, will be required to dissect these mechanical contributions. Future studies will clarify the respective contributions of, and potential feedback between, biochemical signaling and mechanical context (Crozet and Levayer, 2023), and uncover the principles governing the specification and execution of division-type lineages.

While the wing undergoes substantial deformation during pupal development, with overall size staying constant but shape dynamically contracting and elongating, the vein-distance map shows minimal change. Indeed, the terminal position of veins is set with one-cell-width precision (Abouchar et al., 2014). These observations indicate that veins provide robust positional information for coordinating cell-division patterns across a deforming tissue. Because the vein-distance map can itself be altered by cell divisions, feedback between vein positioning and division dynamics may help stabilize and refine tissue morphogenesis.

The initial positions of local chains of division are biased with respect to vein distance, and division events then propagate to neighboring cells. Whether the “seeding” and “spreading” processes are governed by the same signaling pathway, particularly EGFR/ERK signaling, remains an open question. The finding that spatially propagating ERK activity promotes G2/M progression in mammalian epidermis (Hiratsuka et al., 2015) is intriguing; however, our observation that local propagation of cell division also occurs in the veinless wing suggests that EGFR/ERK signaling may not be required for this spreading process or at least is not its sole regulator.

As discussed above, the presence of distinct cell groups with different division dynamics can help preserve tissue integrity. However, if divisions were patterned strictly according to vein distance, such a mechanism would inherently generate division domains in which mechanical stress becomes concentrated. Our numerical simulations, though simple and qualitative, suggest that the observed loosening of this coupling may act to counterbalance the mechanical instability introduced by a vein-distance-dependent control, reducing stress accumulation and helping to prevent deleterious outcomes such as tissue buckling or mechanical cell competition (Tozluoǧlu and Mao, 2021; Matamoro-Vidal and Levayer, 2019). Future quantitative analyses integrating experimentally measured forces and cell dynamics with numerical simulations will be required to further test this possibility.

Finally, our findings raise broader questions about how cell-size control via distinct division types adapts to functional demands and diverse wing architectures. Cell size is a fundamental biological parameter that influences both the efficiency of biochemical reactions and the mechanical properties of cellular structures (Ginzberg et al., 2015). In the *Drosophila* wing, cell-size homogeneity has been proposed to facilitate hexagonal cell packing, which is reciprocally coupled to the PCP alignment (Chhajed et al., 2025; Classen et al., 2005; Ma et al., 2008; Sugimura and Ishihara, 2013). Cell size may also affect insect flight performance by tuning the density and spatial distribution of wing hairs, thereby influencing aerodynamics (Polet et al., 2015). Wing size, venation patterns, and margin shapes vary widely among insect species (Grodnitsky, 1999; Shimmi et al., 2014; Matamoro-Vidal et al., 2015), raising the question of whether the division control identified in *Drosophila melanogaster* are conserved or evolutionally modified to accommodate different wing geometries and patterning. For example, how vein-distance-dependent mechanisms scale with wing size represent interesting directions for future evo-devo studies. In conclusion, through a comprehensive description of cell division and cell-size dynamics, this study provides a framework for addressing important yet unresolved questions about how cell division is coordinated in space and time to support robust morphogenesis and functional tissue design in insect wing epithelia.

## Materials and Methods

### Drosophila genetics

*Drosophila* stocks were maintained at 25℃ or 17℃ in a plastic vial on food made from cornmeal, yeast, sucrose, wheat germ, agarose, propionic acid, Butyl *p*-Hydroxybenzoate, and water (Ikawa et al., 2023). The flies used in the present study were *DE-cad-GFP* (Huang et al., 2009), *rhomboid^ve^* (*rho^ve^*), *vein^1^*(*vn^1^*) (BDSC # 603542), *MS1096-Gal4* (BDSC #8860), *UAS-ds dsRNA* (BDSC # 28008), *UAS-argos^30-85-1^* (segregated from BDSC #5363), and *Cnn-GFP* (Megraw et al., 2002). The fly genotypes were *DE-cad-GFP* (wild type), *DE-cad-GFP, Cnn-GFP* (wild type), *DE-cad-GFP*; *rho^ve^, vn^1^* (veinless), *MS1096/+ or Y*; *DE-cad-GFP/DE-cad-GFP*; *UAS-ds dsRNA/+* (*ds* RNAi), and *MS1096/+ or Y*; *DE-cad-GFP/DE-cad-GFP*; *UAS-argos^30-^*^85-1^*/+* (*argos* overexpression).

### Image collection

Preparation of *Drosophila* pupal wing samples for imaging was performed as previously described (Guirao et al., 2015; Ikawa and Sugimura, 2018). White pupae were collected, with the timing of pupation (*i.e.*, 0 h APF) determined to within 10 minutes. Pupae at the appropriate developmental stages were dissected to remove the pupal case over the left wing and then placed on a small drop of Immersol W 2010 (Zeiss 444969-0000-000) in a glass-bottom dish. Imaging was conducted using an inverted confocal spinning disk microscope (Olympus IX83 with Yokogawa CSU-W1) equipped with an iXon3 888 EMCCD camera (Andor), an Olympus 60×/NA1.2 SPlanApo water-immersion objective, and a temperature control chamber (TOKAI HIT). Data acquisition was carried out using iQ2 or iQ3 software (Andor). Whole-wing DE-cad-GFP images were captured at six or eight positions every 5 min, starting no later than 15.5 h APF, and the images were stitched post-acquisition. Cnn-GFP images were acquired every 1 min starting from 18 h APF.

### Image processing

#### Image segmentation

Images of wild-type and *ds* RNAi wings were segmented using custom-made macros and plugins in ImageJ/Fiji (Ishihara and Sugimura, 2012; Guirao et al., 2015). Images of veinless wings were segmented using **EpySeg** (Aigouy et al., 2020). Manual correction was applied where necessary. Vertex positions and connectivity were extracted from skeletonized images using custom-made code in OpenCV for wild-type and *ds* RNAi wings (Ishihara and Sugimura, 2012), or the **GetVertex** plugin in ImageJ/Fiji for veinless wings (see Data and resource availability). Both codes produce identical outputs.

#### Cell Tracking

Cell correspondences between successive frames were obtained by tracking cells in time-lapse data (Namba et al., 2025). Specifically, displacements between sequential images were estimated using **PIVlab** in MATLAB with FFT window deformation and ensemble correlation. Cells were identified by labeling segmented regions, and tracking was performed by adjusting cell centers with displacement vectors and matching IDs between frames using a cost matrix optimized with **assignjv** in MATLAB. Cell division was identified by the appearance of new cells and the area reduction of neighboring cells, with the greatest reduction marking the dividing cell. Tracking results were manually reviewed for accuracy. In addition, tracks were removed from the cell division list when there was ambiguity in identifying daughter cell pairs. Most of the deleted tracks were located around the anterior crossvein (ACV) or near the boundary of the ROI. Cell tracking data for wild-type DE-cad-GFP wings were previously published in Namba et al. (2025), with further manual corrections applied after publication.

### Quantification of cell geometry

Cell geometry, forces, and division were analyzed in Python with **NumPy**, **SciPy**, and **Pandas** modules unless otherwise noted. **Numba** (https://numba.pydata.org/) was used to speed up the calculation when necessary.

#### Input data

The afore-mentioned custom-made code in OpenCV, or equivalently the **GetVertex** plugin in ImageJ/Fiji, provides vertex positions and connectivity, the vertex composition of each junction and cell, and classification of inner and outer vertices, junctions, and cells at a given frame (timepoint). IDs are assigned to all vertices, junctions, and cells. The image coordinate system is defined such that the top-left corner is (0, 0), the *x*-axis increases to the right, and the *y*-axis decreases downward (*i.e.*, *y*-coordinates are zero or negative). For each cell, the vertices are listed in a counterclockwise order.

#### Neighbor cell pairs

A list of first neighboring cells for each cell at a given frame was obtained from the input data described above.

#### Cell area and cell centroid

Cells are represented by a polygon. From the vertex coordinates and composition for each cell, the apical cell area is calculated using:

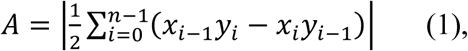

where *n* is the number of vertices of the polygonal cell. *x*- and *y*-coordinates of the cell centroid are given by:

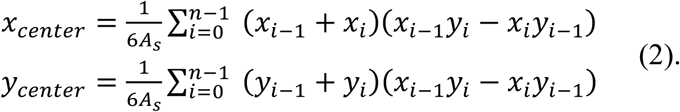

Here, *A_s_* is the signed area of the cell. In Fig. 2E–G, tissue size is defined as the size of the ROI (see below for setting of the ROI) and was calculated by summing the areas of all cells within the ROI.

### Cell division analysis

#### Input data

In our tracking cells in time-lapse data, each track was assigned a unique track ID, which was maintained from the birth to the death of the track. Each track was associated with the corresponding cell ID at each frame, and this information was used as input for the subsequent analysis. In a separate input file, cell division events were listed as follows: frame *t*, track ID of the mother cell at frame *t*, and track IDs of the two daughter cells at frame *t+1*. In this study, the timing of cell division was defined as frame *t+1*.

#### ROI setting

To define the wing-blade ROI, we first manually drew a polygon-shaped ROI boundary at the initial frame (15.5 h APF) and obtained a list of cells that touched the ROI boundary. We then identified cells that persisted until the final frame, regardless of whether they underwent division (permanent lineages). From these permanent lineages, we selected cells that enclosed the imaged area of the wing blade with a polygonal boundary. The same set of permanent lineages was used to define the polygonal ROI throughout all frames, so the ROI boundary was set only once at the beginning. For each frame, we determined whether each cell was located inside this polygon using the Crossing Number Algorithm, and cells inside the polygon were defined as the ROI.

#### Construction of division trees

Division trees were reconstructed by treating each cell track as a node in a rooted binary tree. Division events were stored as tuples (parent, daughter1, daughter2 tracks). Lineages were assembled retrospectively by processing division tuples backward in time and merging daughter nodes into their parent node. After tree construction, graph traversal algorithms were applied to extract lineage structure: breadth-first search was used to enumerate all vertices within each tree, and depth-first search was used to compute node depth and division count. These lineage features were subsequently used to classify division types.

Division trees were reconstructed using data up to 32 h APF in wild-type and *ds* RNAi wings, as retrospective tracking of putative vein cells, identified at 32 h APF, was required (see *Retrospective tracking of putative vein cells* below). For veinless wings, we used data up to 24 h APF, as vein cell tracking was not performed in these samples. As expected from the limited number of cell divisions after 24 h APF (Fig. 1C), the difference in the time period used for lineage reconstruction had minimal impact on the results of this study.

#### Classification of division types

Division events were classified based on the depth of the division tree (Fig. 1B). Divtype 1 refers to a division event that produces two daughter cells from one root cell. Divtype 2 refers to events where granddaughter cells are produced. Divtype 2 is further subdivided into: Divtype 2a, by which one daughter cell and two granddaughter cells are produced and Divtype 2b, by which four granddaughter cells are produced.

Cells belonging to lineages that existed at the initial time frame were used for the analysis of division types. In Fig. 1D and all other figures labeling division types, the following categories of cells were colored gray: cells that did not divide, cells that underwent more than two divisions, cells belonging to lineages excluded from the analysis due to ambiguity in tracking, and cells belonging to lineages that emerged after the initial time frame.

#### Retrospective tracking of putative vein cells

Vein cells can be visually identified at later pupal stages based on their size and shape, brighter DE-cad-GFP signal, and their spatial relationship to sensory cells. We manually labeled a single row of putative vein cells in DE-cad-GFP images at 32 h APF (blue cells in the last frame of Movie 2) and annotated their immediate neighbors (light blue cells in the last frame of Movie 2). These blue and light blue cells were then automatically tracked retrospectively to the beginning of the time-lapse movie. This enabled quantification of the shortest distance from each individual cell to the putative vein cells.

Note that we did not directly monitor the expression of vein or intervein molecular markers during retrospective cell tracking. For this reason, we refer to these retrospectively identified cells as putative vein cells in this manuscript.

#### Detection of local clusters of divided cells

A local cluster of divided cells is defined as a group of six or more neighboring cells that have divided by a given time frame (Fig. 6J, K). These clusters were detected by performing depth-first search-based connected component analysis on the cell-cell neighbor graph. Cells belonging to detected clusters were excluded from further analyses from the subsequent frame onward.

### Bayesian force and stress inference

Force inference methods, in general, solve the inverse problem between cell shape and cellular forces to infer the relative values of junction tensions and pressure differences among cells (Roffay et al., 2021; Sugimura et al., 2016). Bayesian force inference solves the underdetermined problem using Bayesian statistics (Ishihara and Sugimura, 2012).

### Numerical simulations

A Cell Vertex Model (CVM; Fletcher et al., 2014) was used to simulate the progression of cell divisions from veins. The numerical simulations were implemented in C++. Details of the model settings are described in the Supplementary Information.

### Statistics

The data are presented as mean ± SD, computed using the **SciPy** and **NumPy** modules in Python 3.11.8.

## Acknowledgments

The authors would like to thank Yang Hong, Simon Collier, and the Bloomington Stock Center for fly stocks; Kyoko Komano and Miho Aruga for their assistance with image segmentation; Mayu Miyakawa for lab management; Koshi Yoshida for reimplementation of the GetVertex plugin; Tetsuhisa Otani for critical reading of the manuscript; and Yohanns Bellaïche and Lance Davidson for discussion.

## Competing interests

The authors declare no competing interests.

## Author contributions

K.S. designed the research and performed the experiments. K.S., T.N., and S.I. conducted image segmentation, with assistance from Z.Q. T.N. and R.T. implemented the pipeline for cell tracking and division tree construction. K.S., R.T., and T.N. analyzed the data. S.I. performed the numerical simulation. K.S. drafted the manuscript. All authors approved the final manuscript.

## Funding

This study was financially supported by JSPS KAKENHI Grant (24K02029 and 25H01364) to K.S., and JST CREST (JPMJCR1923) and JPJSBP (JPJSJPR 201915010) to S.I. R.T. was supported by the JST SPRING program (JPMJSP2108). Z.Q. was supported by the K. Matsushita Foundation.

## Data and resource availability

The authors declare that the data supporting the findings of this study are included in the paper and its supplementary files. Codes for GetVertex, Bayesian force inference, and CVM simulation can be downloaded from Github: GetVertex (https://github.com/Sugimuralab/GetVertexPlugin), Bayesian force inference (https://github.com/IshiharaLab/BayesianForceInference), and CVM (https://github.com/IshiharaLab/CVM_celldiv_vein). Code for the cell division analysis pipeline will be made available after the related paper by Takayanagi et al. (in preparation) is published. This study did not generate new unique reagents.

## Use of AI tools

ChatGPT (OpenAI) was used for grammar checking, improving readability of the manuscript text, and assisting with code implementation and debugging. The authors reviewed and verified all AI-generated text and code outputs before use in the study.

## Movie legends

**Movie 1 Spatial distribution of cells with different division types in wild-type wing.**

Cells are color-coded according to their division type (divtype). Green: divtype 1; orange: divtype 2; gray: cells with more than two divisions; white: cells that did not divide within the ROI or were located outside the ROI. Colors fade with successive divisions.

**Movie 2 Division events overlaid on vein cells in wild-type wing.**

Cells generated by divisions occurring within the ROI are colored yellow. Putative vein cells are shown in blue and light blue.

**Movie 3 Spatial distribution of cells with different division types in veinless wing.**

Cells are color-coded as in Movie 1.

**Movie 4 Spatial distribution of cells with different division types in *ds* RNAi wing.**

Cells are color-coded as in Movie 1.

**Movie 5 Division events overlaid on vein cells in *ds* RNAi wing.**

Cells are color-coded as in Movie 2.

**Movie 6 Division events in veinless wing.**

Cells generated by divisions occurring within the ROI are colored yellow in the veinless wing.

## SUPPLEMENTARY INFORMATION

## Temporal profiles of cell size during pupal wing development

In Fig. 2A–C, we analyzed the temporal profiles of cell size. Below, we provide supplementary analyses.

### Vein and intervein cells

Retrospective tracking of putative vein cells allowed us to analyze putative vein cells and putative intervein cells separately (Materials and Methods). While vein cells were consistently smaller than intervein cells, both cell types followed a similar temporal pattern in cell size change: their average cell size decreased until around 22 h APF and was then maintained (Fig. S3).

### Cell size changes prior to or following cell division

To directly monitor the cell size changes prior to or post cell divisions, the data from individual cell tracks were aligned in time with respect to their time of cell division (summarized in Fig. 2D; Fig. S4, S5). Tracks lasting less than three frames were excluded from the subsequent analysis because they were likely generated by segmentation and/or tracking errors. In addition, cells enlarge during the mitotic (M) phase, as shown by the last few time points in Fig. S4. To avoid conflating mitotic rounding with interphase growth, data from the M phase were excluded when analyzing post-division size changes of daughter cells that undergo a second round of division (Fig. S5). To do this, we first estimated the M-phase duration as 19.4 ± 2.6 min (n = 60) by measuring the time from the appearance of two Centrosomin (Cnn)-GFP dots to the formation of a nascent junction (Fig. S6). Accordingly, we excluded data from four frames (20–5 min before division) in Fig. S5C, D.

Our analysis revealed distinct temporal profiles of cell size for each division type. Specifically, divtype 1 cells halved their size, from ∼32 µm^2^ to ∼15 µm², through a single round of division (Fig. S4A, S5A). Divtype 2b cells halved their size, from ∼38 µm^2^ to ∼20 µm^2^, at the first division (Fig. S4C, E, S5D). Their size then gradually increased up to ∼25 µm^2^ before being halved again by the second division, followed by a subsequent increase to ∼14 µm^2^ (Fig. S4E, S5D, F). The initial size of divtype 2a cells was ∼33 µm^2^, nearly same as that of divtype 1 cells (Fig. S4B). After the first division, cells that underwent a second division were slightly larger than those that did not, and this difference increased until the second division occurred (Fig. S4D; compare Fig. S5B and Fig. S5C). After the second division, the size profile of divtype 2a cells resembled that of divtype 2b cells (Fig. S5E). Despite these distinct temporal trajectories, the final cell sizes converged toward similar values. In relative order, the largest final size was observed in divtype 2a cells that divided only once, followed by divtype 1 cells, and then by divtype 2b cells and divtype 2a cells with two divisions.

## Estimation of M phase duration

In Fig. S6, we estimated the duration of M phase to be 19.4 ± 2.6 min (n = 60) by measuring the time from the appearance of two Cnn-GFP dots to the formation of a nascent junction labeled with DE-cad-GFP. This measurement was used to determine the excluded time points in Fig. S5C, D.

Here, we discuss this time interval in relation to previous studies. In *Drosophila* and other animal cells, the appearance of Cnn dots corresponds to the assembly of pericentriolar material at the onset of M phase (Conduit et al., 2015). It has been reported that the time between the initiation of chromosome condensation at the onset of M phase and nuclear envelope breakdown (NEBD) is approximately 6 min in *Drosophila* embryos (Fasulo et al., 2012). In *Drosophila* wing disc cells, the interval from NEBD to the completion of spindle formation lasts about 15 min (Bergstralh et al., 2016), based on the sum of two reported time intervals: one from NEBD to the appearance of a complete spindle, and another corresponding to spindle formation itself. Taken together, the total duration of M phase is approximately 21 min, indicating that our measurement using Cnn-GFP and DE-cad-GFP is consistent with previous studies.

## Discussion on cell division progression from wing margins

In addition to the division waves correlated with the distance from veins, we observed that cell divisions also initiated near the wing margins, particularly along the posterior sides of the wing (Fig. 3A). This observation is consistent with the accumulation of *string* (*cdc25*) transcript-positive cells along the wing margins at 12–16 h APF (Milán et al., 1996). In wild-type pupal wing movies acquired by others and by us, however, the distal-most margins were not captured at early developmental stages (Aigouy et al., 2010; this study), and therefore potential distal margin-initiated divisions could not be directly assessed. By contrast, the geometry of the veinless wing allowed visualization of the distal margin from the beginning of the movie, revealing two directed waves of divisions emerging from both proximal and distal sides (Fig. 4D). Whether a comparable distal-originating wave exists under wild-type conditions will require further investigation. Nevertheless, even if a distal wave were present in wild type, its contribution would be significantly smaller than in the veinless mutant. In wild type, the division wave originating from the proximal side traverses the central region and reaches the distal side. This contrasts with the veinless mutant, in which a distal-originating wave progresses into the central region.

## Discussion on veinless mutant phenotypes

In this study, we examined the influence of veins by perturbing EGFR signaling and, consequently, vein fate, using the veinless mutant (De Celis, 1998; Shilo, 2005). Because EGFR signaling is known to promote cell proliferation, the observed phenotypes in the veinless mutant could in principle be attributed to a general reduction in cell division caused by perturbed EGFR signaling. However, as shown in Fig. 4B, the veinless mutation selectively suppresses the divtype 2b lineage while leaving the divtype 1 lineage largely intact. Such selective suppression of a specific lineage is difficult to reconcile with a general inhibition of cell proliferation. Moreover, observations in wild-type wings further support the idea that division-type specification is linked to vein-associated spatial context: in the region near the PCV, which is specified later (Blair, 2007; Matsuda and Shimmi, 2012), division progression is delayed, and fewer divtype 2 cells are observed in intervein cells abutting the PCV compared with those abutting L4 or L5 (arrows in Fig. 3A, B). In line with this, discontinuity of the L4 vein upon overexpression of Argos, a negative regulator of EGFR signaling (Blair, 2007), results in reduced cell division in the vicinity of L4 (Fig. S9). Therefore, the results from wild-type and genetically modified wings suggest that division dynamics including division-type specification are influenced by vein-associated spatial cues.

In the veinless wing sample shown in the main figures, the remaining divtype 2 cells are concentrated in the proximal-posterior region of the wing (Fig. 4C). Although weaker, a similar tendency is observed in another veinless sample. Their spatial distribution does not fully coincide with the expected position of L5, suggesting that residual vein identity is unlikely to fully account for this pattern. Instead, this enrichment could potentially reflect positional cues associated with the posterior wing margin. Further analyses will be required to determine the origin and regulation of these remaining divtype 2 cells.

In the veinless wings, the probability of undergoing apoptosis is increased to 1.92 ± 0.06-fold relative to the mean in wild-type wings (n = 3 wild-type and n = 2 veinless). Importantly, the deviation in the tissue size–progeny number relationship (Fig. 2G) is observed even though apoptotic cells are not included in the progeny counts. This indicates that altered coordination of division types, rather than increased apoptosis, plays a substantial role in the misregulation of cell size in the mutant.

## Numerical simulations

We performed numerical simulations with scheduled and triggered cell-state transitions. The simulations use a two-dimensional Cell Vertex Model (Fletcher et al., 2014) consisting of vein cells (V cells) and other wing cells (hereafter referred to as wing cells). For the sake of simplicity, we set V cells as non-proliferative. Wing cells undergo one round of cell division through the state sequence I → M → D, where I, M, and D denote initial, mitotic, and divided states.

### Cell vertex model

In the Cell Vertex Model, the cell geometry is represented as a tiling of polygonal cells. The mechanical energy is defined as

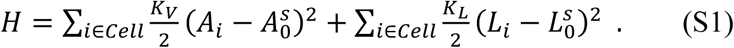

Here, *A*_i_ is the area of the *i*-th cell, *L*_i_ is the contour length of the *i*-th cell, and the preferred area 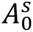 and preferred contour length 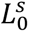 depend on the cell type and state *s* ∈ {*V*, *I*, *M*, *D*} (Table A). The shape index *ϕ*, defined as the ratio of the preferred contour length to the square root of the preferred cell area, was fixed at *ϕ* = 3.7, for which a hexagonal cell shape is favored (Bi et al., 2016). Changes in tissue geometry are governed by the time evolution of the vertex position ***r***_j_, given by

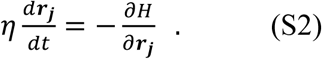

We numerically solved Eq. S2 while allowing vertex reconnections, corresponding to T1 transitions. The initial condition was prepared by relaxing a system composed of V and I cells for a certain period of time, with I cells occupying the region between the V cells (First frame in Fig. S12A–C). The initial numbers of V and I cells were 199 and 301, respectively, giving a total initial cell number of *N*_0_ = 500. Equation (S2) was numerically solved under periodic boundary conditions using the second-order Runge–Kutta algorithm with Δ*t* = 1.0 × 10^/0^. One simulation time unit corresponds to 2 min, and the simulations were performed over 0–500 min. The parameter values are listed in Table B and were chosen so that the simulated area change during the mitotic phase was within a reasonable range.

#### Cell state transitions

Wing cells first transition from the I-state to the M-state, accompanied by a 1.5-fold increase in preferred size, 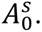 After 20 min, each M cell undergoes division with a random orientation and generates two D cells, whose preferred size is set to half that of an I cell (Table B). The I → M transitions follow Scenario A, B, or C, which differ in the rule determining the timing of the I → M transition (Fig. S12A–C). In these scenarios, transitions occur either spontaneously (scenarios A and B) or deterministically (scenario C) or are induced by neighbors (scenarios A and B). In the latter case, an I cell has a higher probability of transitioning into the M-state when it has neighboring M or D cells.

##### Scenario A

The first division events are initiated only in cells adjacent to veins, and divisions subsequently spread through the neighbor effect (Fig. S12A).

i. An I cell adjacent to a V cell transitions into an M cell at a rate of 0.1 min⁻¹ (spontaneous transition). This ensures that most vein-adjacent cells transition into the M-state during the early stage, within approximately 10 min.
ii. Other I cells that are not adjacent to any V cells transitions into an M cell only when they have neighboring M or D cells. The transition probability is given by 1 − (1 − *r*Δ*t*)^1^ in each time step, where *r* = 0.05 min⁻¹ and *n* denotes the number of neighboring M and D cells.

These rules lead to stochastic progression of cell division from the vein.

##### Scenario B

Division events are initiated by vein-distance-dependent spontaneous seeding and spread through the neighbor effect (Fig. S12B; motivated by the experimental observations in Fig. 6H–K).

i. At *t* = 0, each I cell is assigned a scheduled spontaneous transition time drawn from an exponential distribution with a position-dependent mean. The mean depends on the positional coordinate *y* of the cell centroid, measured along the vertical axis, such that cells closer to veins have earlier scheduled transition times. Specifically, it is given by *τ*(*y*) = 1,600 × *f*(*y*) min, where *f*(*y*) is a piecewise linear function satisfying *f*(0) = *f*(*L*) = 0 and *f*(*L*/2) = 1. Here, *L* denotes the system length along the vertical axis (Table B). An I cell spontaneously transitions into the M-state once its scheduled transition time is reached.
ii. If an I cell has neighboring M or D cells, the transition to the M-state is triggered even before its scheduled transition time. The transition probability in each time step is set in the same manner as in Scenario A: 1 − (1 − *r*Δ*t*)^n^, where *r* = 0.05 min^-1^ and *n* denotes the number of neighboring M and D cells.

These rules lead to the combination of stochastic spontaneous transition and progression of cell division from existing M or D cells. With these parameter values, the total number of spontaneous cell divisions is nearly identical between Scenarios A and B.

##### Scenario C

Progression of cell divisions is deterministically controlled by vein distance, without neighbor-dependent spreading (Fig. S12C).

i. An I cell at position *y* transitions into an M cell at time 150 × *f*(*y*).

This rule produces deterministic progression of cell divisions according to the distance from the vein. The parameter value was chosen such that the division rate in Scenario C was comparable to the approximately constant division rate observed during the intermediate phase of Scenario B. As shown in Fig. S12F, the resulting progression time in Scenario C is slightly shorter than that in Scenario B, but remains comparable.

---

**Figure S1.**
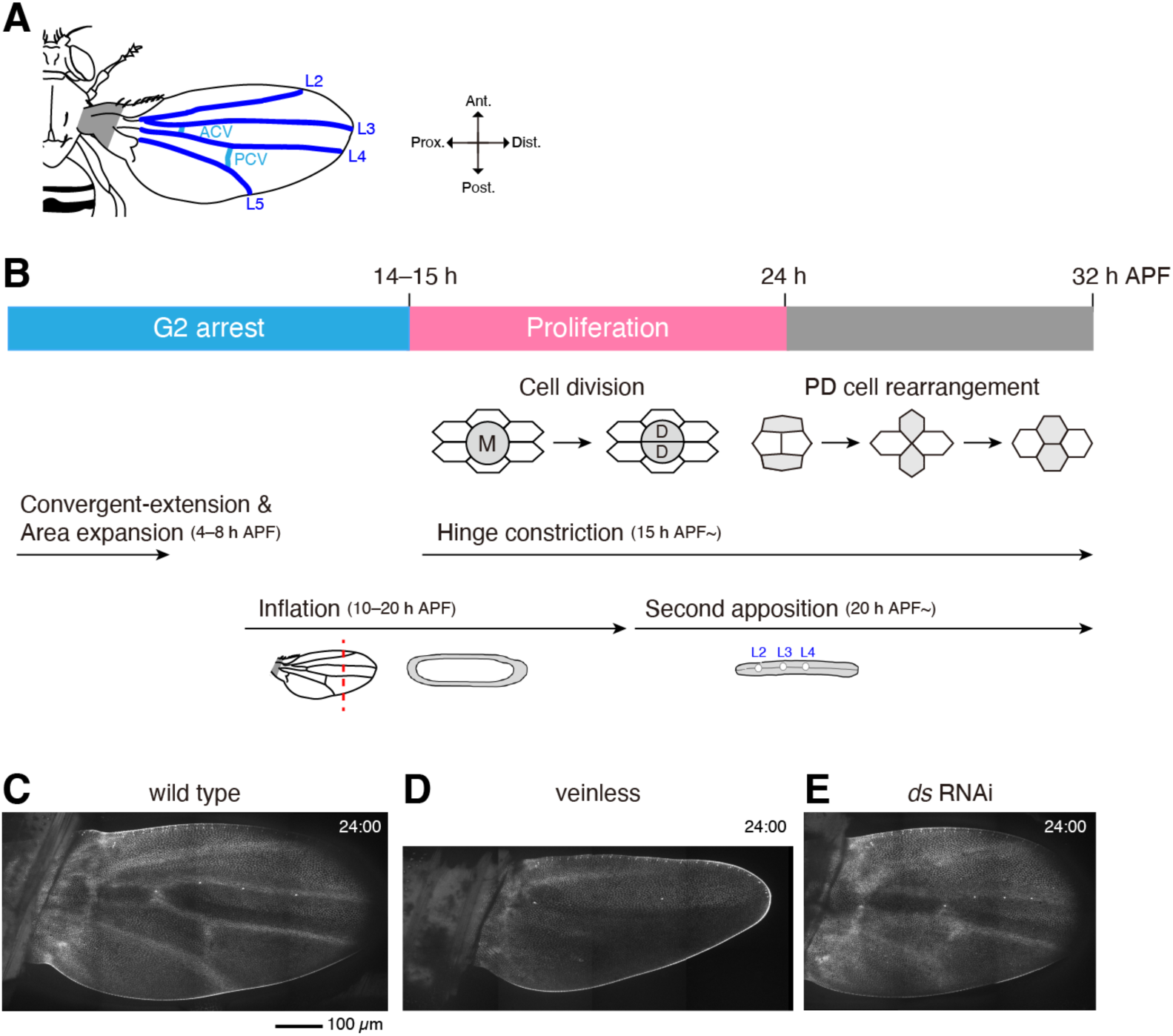
Time course of *Drosophila* pupal wing development. (A) Schematic of *Drosophila* adult wing. Longitudinal veins (L2–L5) are shown in blue, and cross veins, anterior cross vein (ACV) and posterior cross vein (PCV), are shown in light blue. The hinge region is shaded in gray. In this and all subsequent figures, the vertical and horizontal axes correspond to the anterior-posterior (AP) and proximal-distal (PD) axes, respectively. (B) Time course of pupal wing development is illustrated based on the observations of previous studies (see the Introduction for references). During early pupal stages, wing cells are arrested in the G2 phase. As ecdysone signaling declines, cells resume the cell cycle and begin to proliferate. Most cell divisions cease by 24 h APF. Hinge constriction begins at ∼15 h APF, generating PD-oriented tissue tension. Subsequently, PD-oriented cell intercalation occurs from ∼21–22 h APF as a passive response to this uniaxial stretching. Although not analyzed in this study, starting from 4 h APF the wing epithelium undergoes convergent extension to elongate along the PD axis. This is followed by a phase of isotropic expansion, which increases apical cell area (Diaz-de-la-Loza et al., 2018). Bottom panels show lateral views of the wing along the red dotted line during the inflation and second apposition stages. After the first apposition stage (not shown), the dorsal and ventral epithelial layers separate between 10–20 h APF (inflation) (Blair 2007; Montanari et al. 2022). The two layers then reattach at ∼20 h APF (second apposition), with a small gap remaining along the veins. (C–E) Images of pupal wing expressing DE-cad-GFP at 24 h APF. Wild type (C); veinless (D); *ds* RNAi (E). (A) and (B) are adapted from Ikawa and Sugimura (2018). Schematics of wing lateral views in (B) are adapted from Montanari et al. (2022). Scale bar: 100 µm (C).

**Figure S2.**
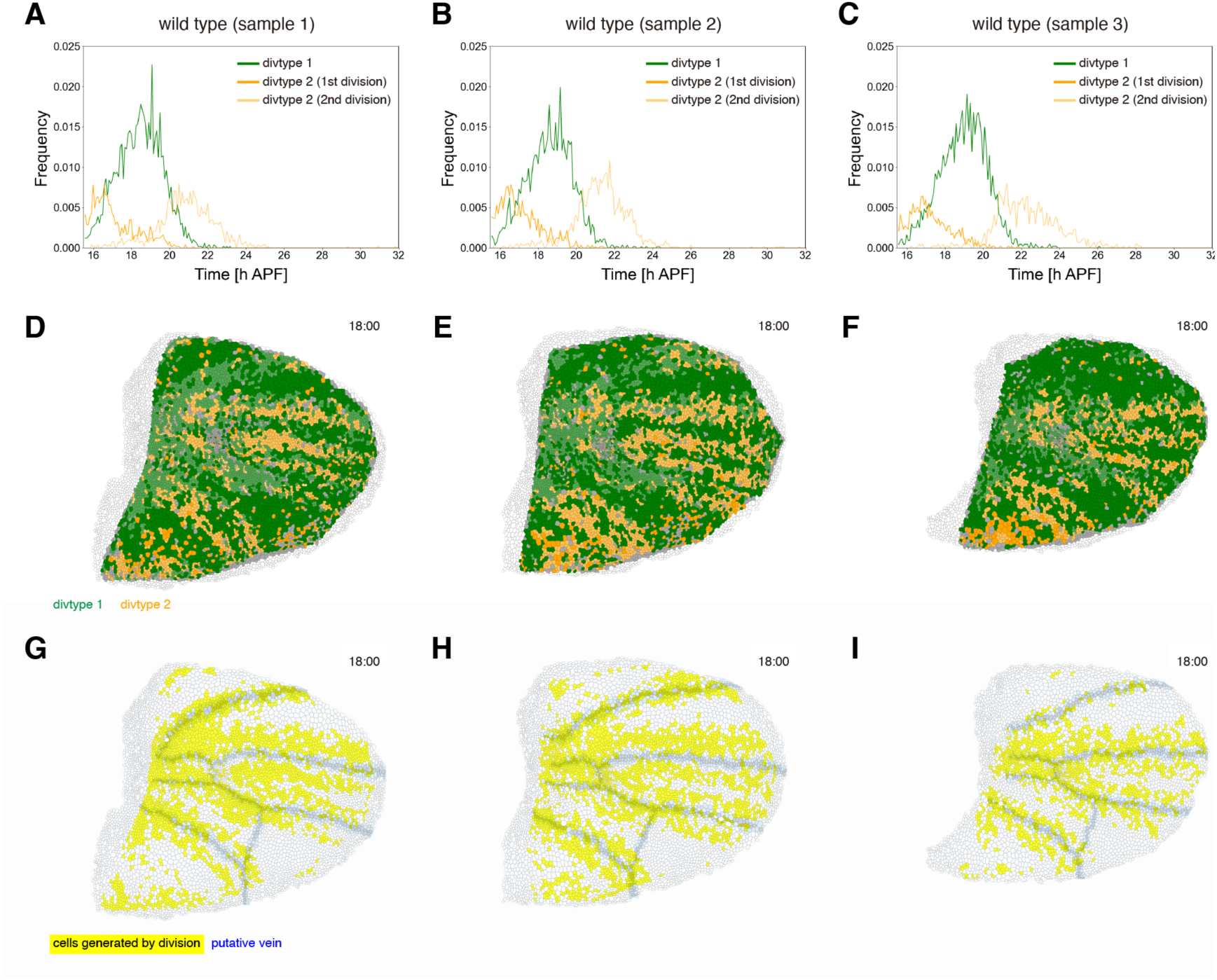
Reproducibility of division timing and positions within the wing. (A–C) Frequencies of each divtype are plotted over developmental time for individual samples (green: divtype 1; orange: first division in divtype 2; light orange: second division in divtype 2). (D–F) Cells belonging to lineages that existed at the initial time point are color-coded according to their division type. Snapshots are shown at 18 h APF. Green, divtype 1; orange, divtype 2; gray, other cells within the ROI; white, cells outside the ROI. Colors fade with successive divisions. (G–I) Cells generated by divisions occurring within the ROI are colored yellow at 18 h APF. Putative vein cells are shown in blue or light blue.

**Figure S3.**
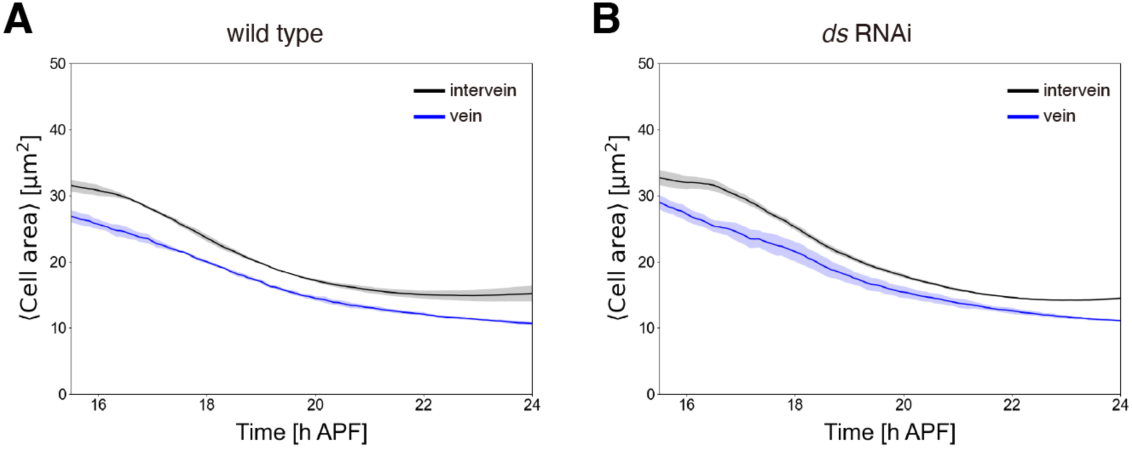
Temporal profiles of cell area for putative vein and intervein cells. (A, B) Time profiles of average area for putative intervein cells (black) and putative vein cells (blue). Wild type (A); *ds* RNAi (B). Data are plotted as mean ± SD. n = 3 wild-type wings (A). n = 3 *ds* RNAi wings (B).

**Figure S4.**
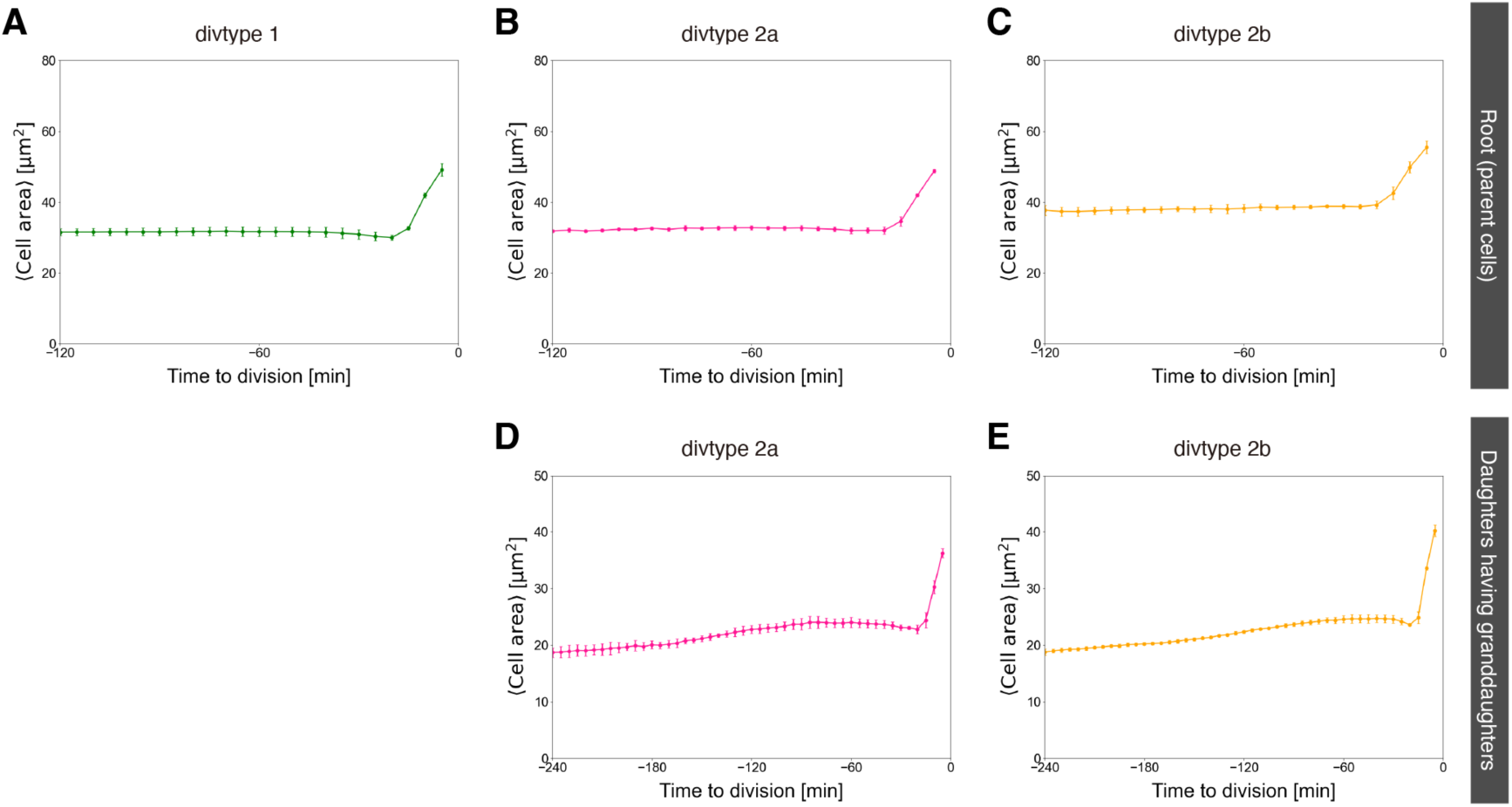
Temporal profiles of cell area prior to cell division. (A–C) Cell area was quantified for each cell track prior to its division. (A) Root cells of divtype1. (B) Root cells of divtype 2a. (C) Root cells of divtype 2b. Timepoint *t* = 0 corresponds to the timing of cell division, when a single cell divides into two. An increase in cell area in the last few time points corresponds to mitotic rounding. (D, E) Same analysis as in (A–C), applied to the second layer of the division tree. (D) Daughter cells of divtype 2a that underwent subsequent division. (E) Daughter cells of divtype 2b. Data are plotted as mean ± SD. n = 3 wild-type wings (A–E).

**Figure S5.**
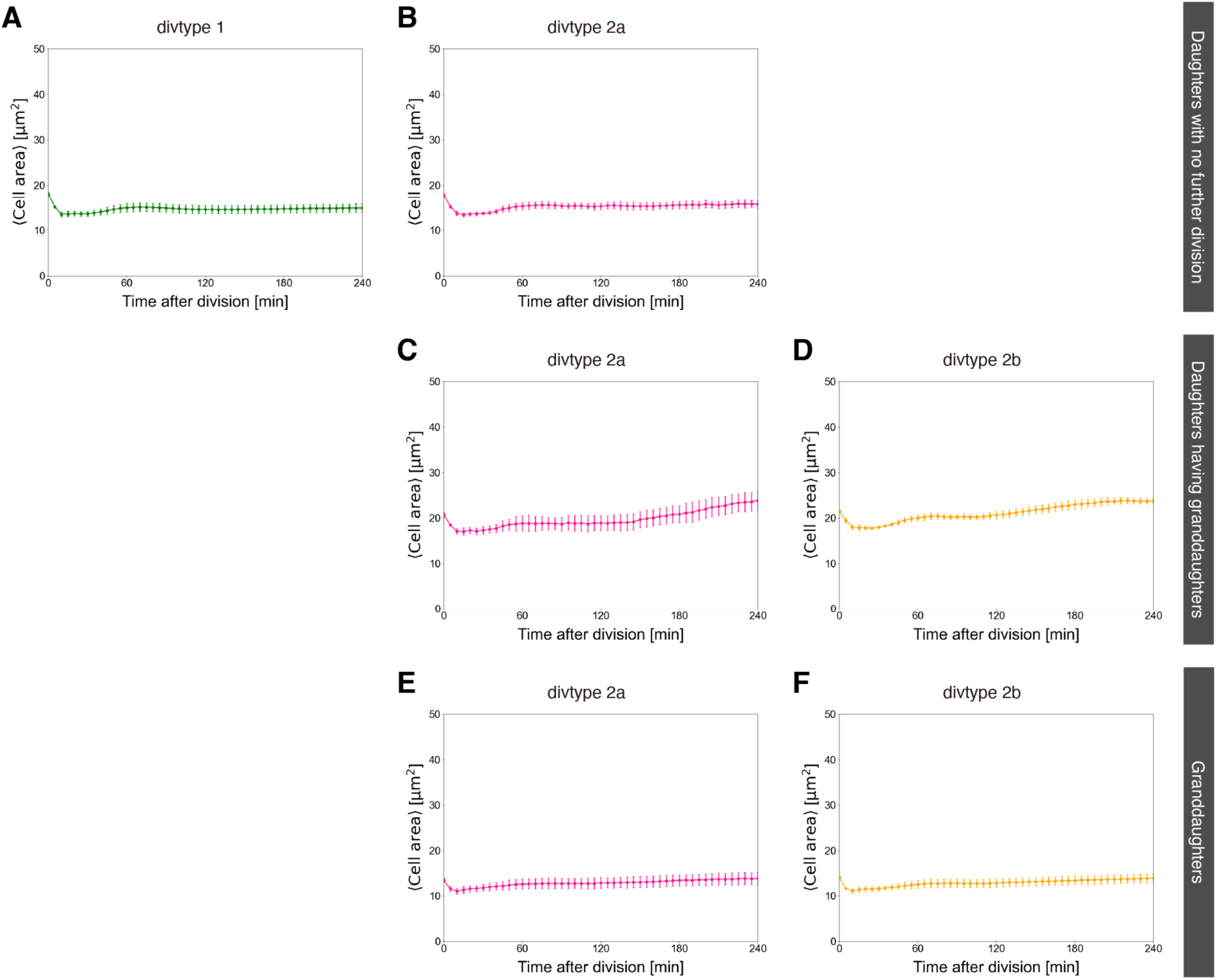
Time profiles of cell area following cell division. (A–D) Cell area was quantified for each cell track after a preceding division event. (A) Daughter cells of divtype 1. (B) Terminal daughter cells of divtype 2a (*i.e.*, cells that did not divide again). (C) Daughter cells of divtype 2a that underwent subsequent division. (D) Daughter cells of divtype 2b. Timepoint *t* = 0 corresponds to the timing of cell division, when a single cell divides into two. We excluded data from four frames (20–5 min before division) in (C) and (D) (see Supplementary Information and Fig. S6 for the estimation of the M phase duration). (E, F) Same analysis as (A–D), applied to the third layer of the division tree. (E) Granddaughter cells of divtype 2a. (F) Granddaughter cells of divtype 2b. Data are plotted as mean ± SD. n = 3 wild-type wings (A–F).

**Figure S6.**
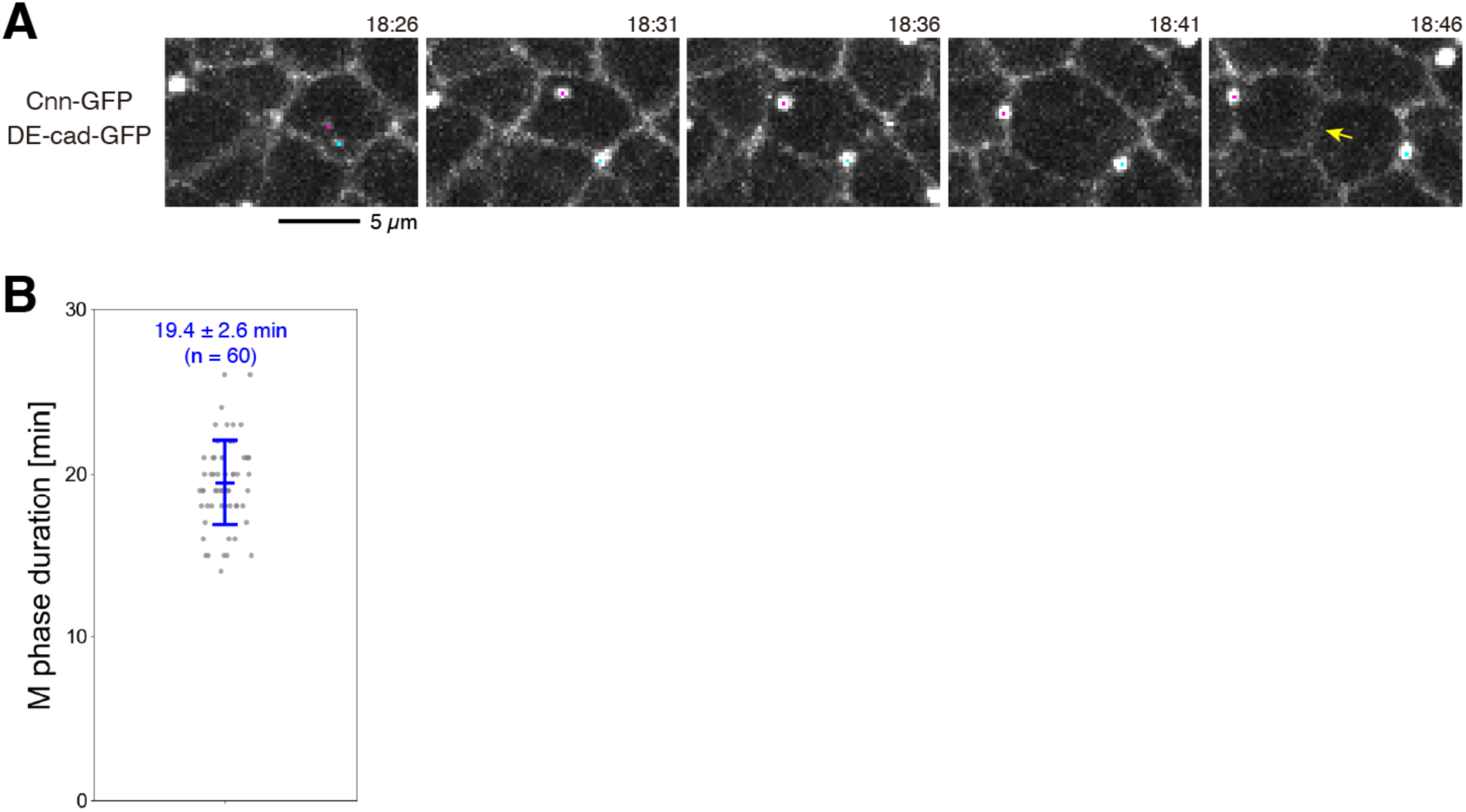
Quantification of the M phase duration in pupal wing. (A) Time-lapse images of wing cells expressing DE-cad-GFP and Cnn-GFP. Magenta and blue dots indicate centrosomes. Yellow arrow marks the formation of a nascent junction following cell division. (B) From time-lapse images of DE-cad-GFP and Cnn-GFP, the duration from the appearance of two Cnn-GFP dots to the formation of a nascent junction was quantified and plotted. Gray dots represent the data from each division. The number of divisions analyzed is indicated. Data are plotted as mean ± SD (B). Scale bar: 5 µm (A).

**Figure S7.**
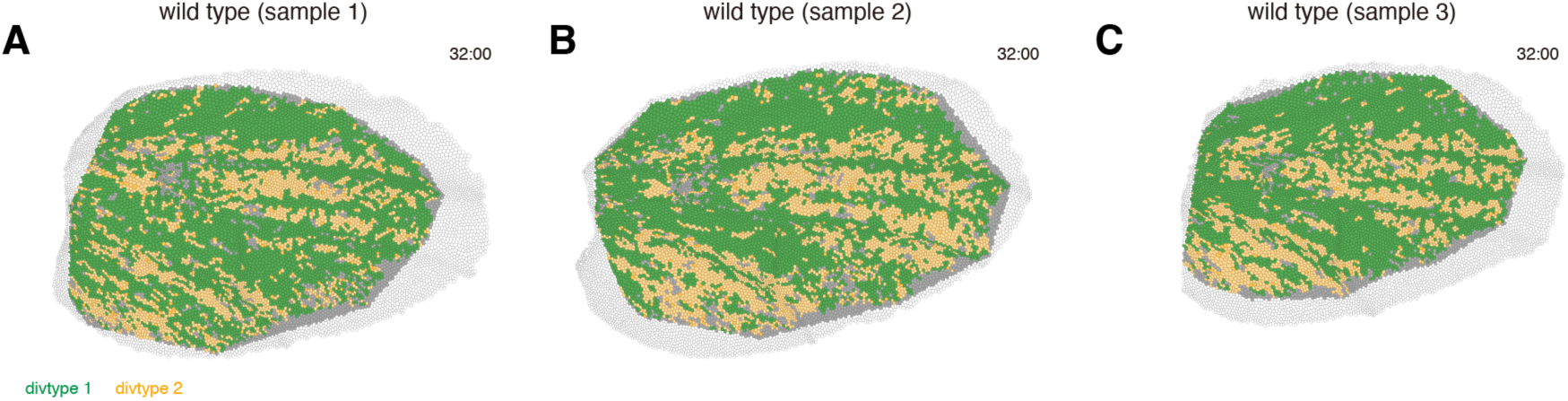
Spatial arrangement of different divtype cells at 32 h APF. (A–C) Wild-type wing cells belonging to lineages that existed at the initial time point are color-coded according to their division type. Snapshots are shown at 32 h APF when the PD cell rearrangement process are complete. Green, divtype 1; orange, divtype 2; gray, other cells within the ROI; white, cells outside the ROI. Colors fade with successive divisions.

**Figure S8.**
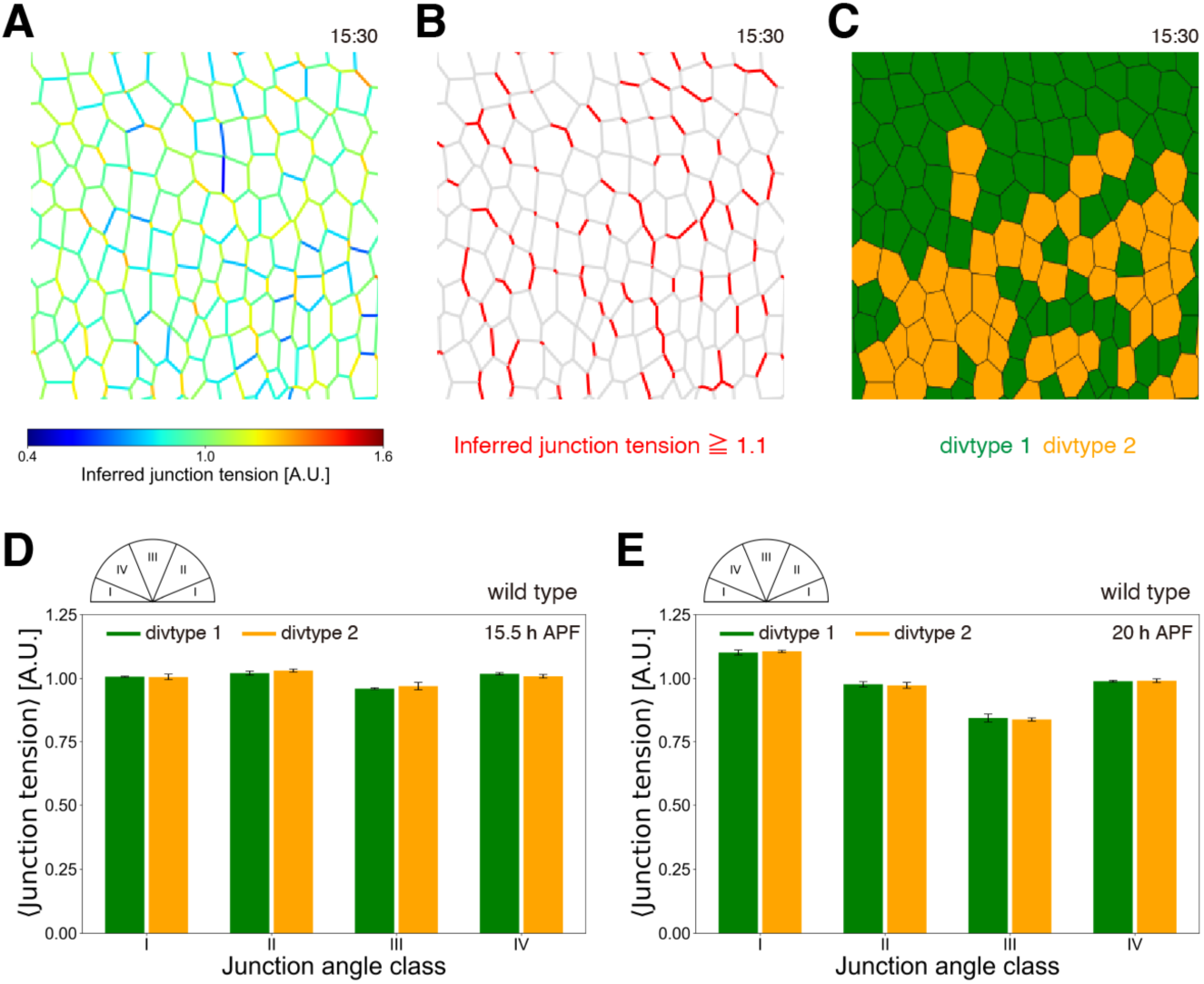
No significant correlation between tension anisotropy and division type. (A–C) Cells located between the L2 and L3 veins in a wild-type wing at 15.5 h APF. (A) Map of junction tension inferred by Bayesian force inference (Ishihara and Sugimura, 2012; Materials and Methods). The color bar indicates the relative values of junction tension. (B) Junctions with inferred tension ≥ 1.1 are colored red, and all other junctions are colored gray. (C) Division types are colored as in Fig. 1D. (D, E) Mean junction tension for each junction-angle class at 15.5 h APF (D) and 20 h APF (E) for cells in divtype 1 and divtype 2 lineages (green and orange, respectively). Junction-angle classes are defined as shown in the semicircle at the top left. Data are plotted as mean ± SD (D, E). n = 3 wild-type wings (D, E).

**Figure S9.**
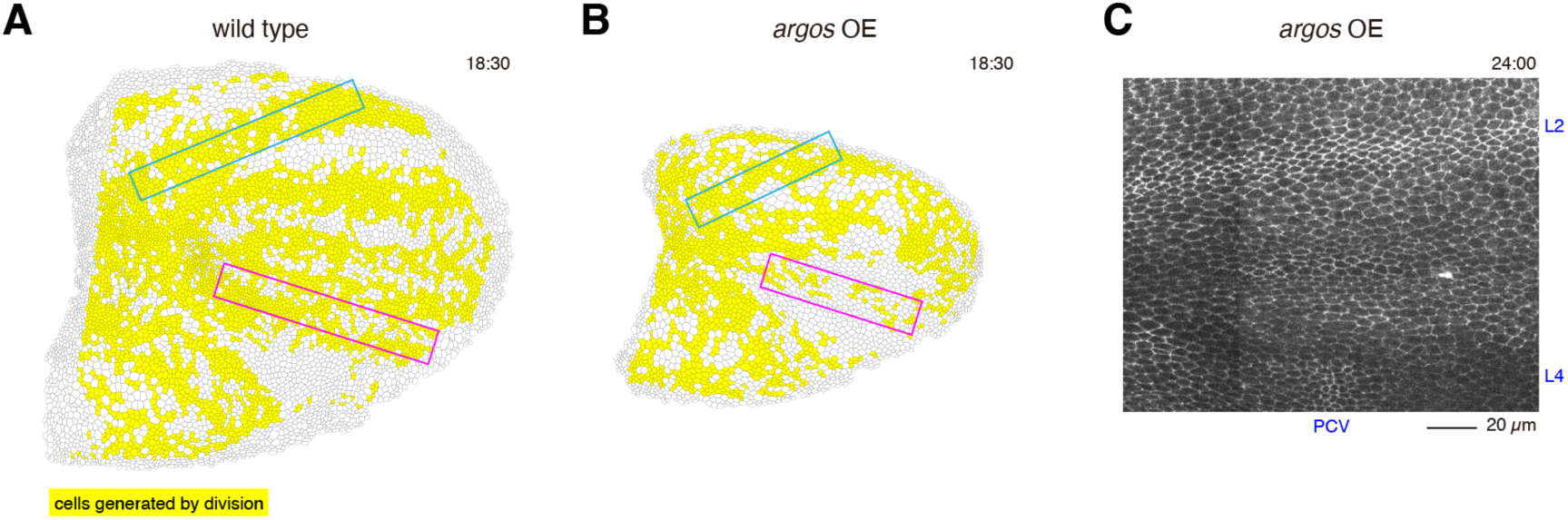
Disruption of the L4 vein is accompanied by reduced cell division. (A, B) Cells generated by divisions within the ROI are colored yellow at 18.5 h APF (A: wild type; B: *argos* overexpression (OE)). Cell divisions around L4 are less frequent in *argos* OE wings than in wild type. Blue and magenta rectangles indicate regions around the L2 and L4 veins, respectively. (C) Magnified images of DE-cad-GFP at 24 h APF from the same *argos* OE wing shown in (B). The L4 vein is largely absent, whereas the L2 vein remains intact. Similar vein phenotypes are also observed in adult wings. Scale bar: 20 µm (C).

**Figure S10.**
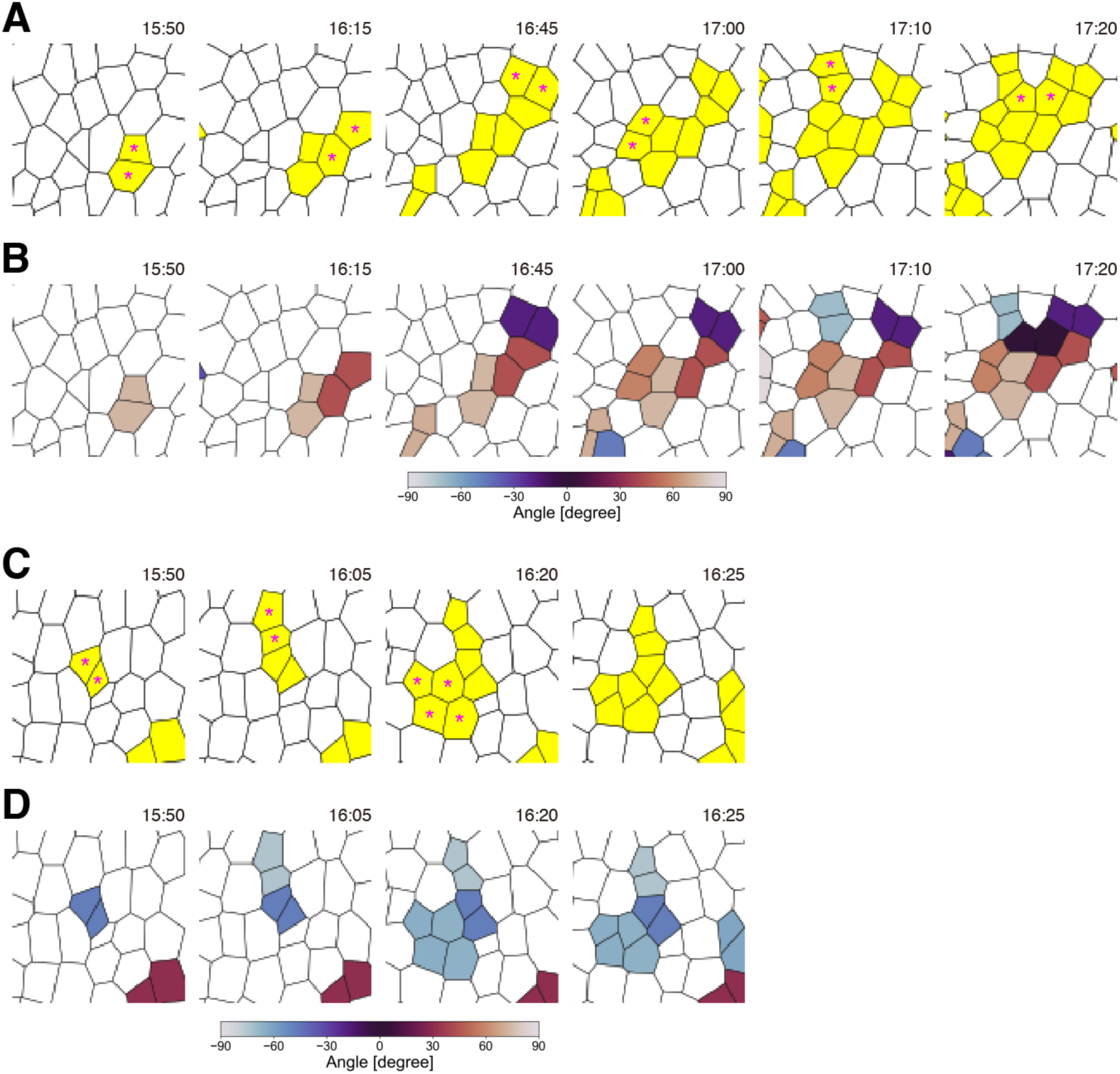
Additional examples of local progression of cell division events. (A) Cells generated by division are shown in yellow, and newly divided cells in the current frame are marked with asterisks. Developmental time is indicated in the top-right corner. (B) Cells generated by divisions are color-coded according to their division orientation (see color bar at the bottom). (C, D) Same as (A) and (B), but from a different wild-type sample.

**Figure S11.**
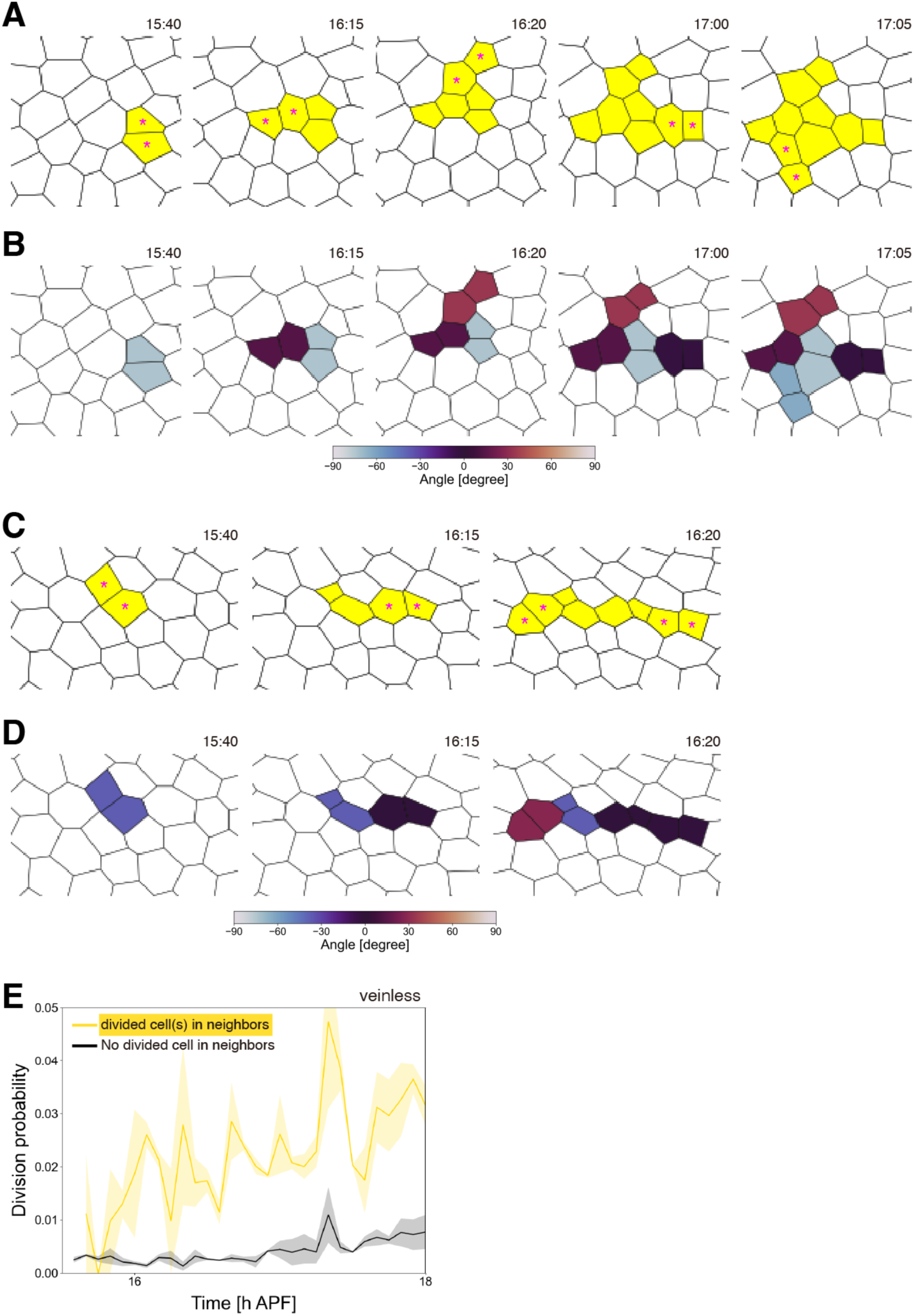
Local progression of cell division events in veinless wing. (A, C) Cells generated by division are shown in yellow, and newly divided cells in the current frame are marked with asterisks, as in Fig. 6D, F and Fig. S10A, C. (B, D) Cells generated by division are color-coded according to their division orientation, as in Fig. 6E, G and Fig. S9B, D. (E) Division probability is compared in the presence and absence of divided neighbors (see the legend of Fig. 6H for details). Yellow lines represent cells with at least one divided neighbor; black lines represent cells with no divided neighbors. Data are plotted as mean ± SD (E). n = 2 veinless wings (E).

**Figure S12.**
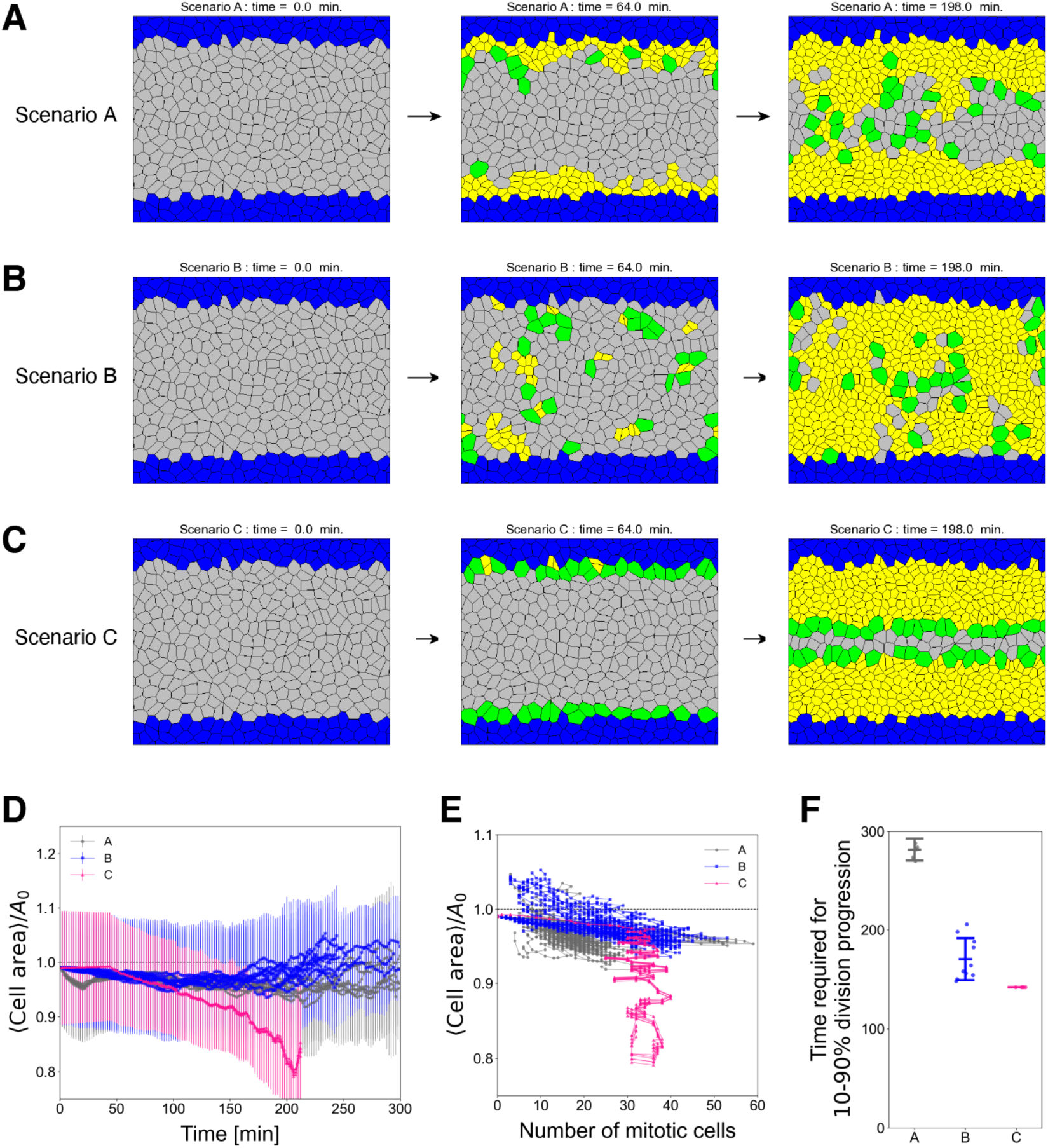
Numerical simulations of cell division progression. (A–C) Snapshots from simulations based on different rules governing cell division progression (A: scenario A; B: scenario B; C: scenario C; see Supplementary Information for details). Vein cells (blue) are positioned at the top and bottom boundaries of the system. For the sake of simplicity, vein cells are treated as non-proliferative. Other wing cells undergo one round of cell division through the state sequence I (initial; gray) → M (mitotic; green) → D (divided; yellow). In (A), the first division events are initiated only in cells adjacent to veins. Subsequently, the transition to the M-state is triggered when I cells have neighboring M or D cells. In (B), each I cell is assigned a scheduled spontaneous transition time based on its distance from veins along the vertical axis. An I cell spontaneously transitions into the M-state once its scheduled transition time is reached. The transition to the M-state is also induced by neighbor effects, as in (A). These settings are motivated by the experimentally observed combined effects of vein-dependent seeding and neighbor-dependent spreading (Fig. 6H–K). In (C), the timing of transition to the M-state is strictly determined by vein distance, without neighbor effects. This deterministic control is reflected in the nearly identical trajectories in (D) and (E) and the zero standard deviation in (F). To reliably compare results among different scenarios, parameter values were chosen such that (1) the total number of spontaneous cell divisions was nearly identical between Scenarios A and B, and (2) the division rate in Scenario C was comparable to the approximately constant division rate observed during the intermediate phase of Scenario B. (D) Time evolution of the mean cell area for three scenarios (gray, scenario A; blue. scenario B; magenta, scenario C). The mean cell area is normalized by a target cell area 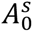 listed in Table A. Data are plotted until the time point at which 95% of I cells have completed the transition to the M-state. (E) The normalized mean cell area is plotted against the number of mitotic cells at each time point. Colors represent different scenarios (gray, scenario A; blue, scenario B; magenta, scenario C). Data are plotted until the time point at which 95% of I cells have completed the transition to the M-state. (F) Time required for division progression in scenarios A, B, and C. The elapsed time from 10% to 90% of wing cells undergoing division is quantified and plotted. Dots represent data from individual simulation runs. Data are plotted as mean ± SD (D, F). The number of simulation runs were 7, 10, and 6 for scenario A, B, and C, respectively (D–F).

**Table A.** Cell type- and state-specific parameters. Wing cells undergo the state transition sequence I → M → D and divide once. In all scenarios, after an I cell transitions into the M-state, it subsequently transitions into the D-state after 20 min.

| Symbol | Meaning | Preferred area | Preferred contour |
| --- | --- | --- | --- |
| V | Vein cell; no cell division<br>Parameters do not change | $A_0^V = A_0$ | $L_0^V = L_0$ |
| I | Initial state of other wing cells | $A_0^I = A_0$ | $L_0^I = L_0$ |
| M | Mitotic phase (duration: 20 min)<br>Preferred size is increased by 1.5-fold | $A_0^M = 1.5A_0$ | $L_0^M = \sqrt{1.5}L_0$ |
| D | Divided state<br>Preferred size is reduced to half | $A_0^D = A_0/2$ | $L_0^D = L_0/\sqrt{1.5}$ |

**Table B.** Other parameters used in numerical simulations. The table lists baseline parameters and simulation settings.

| Quantity | Value / Definition |
| --- | --- |
| System size | $L \times L = 1.0 \times 1.0$ |
| Initial number of cells | $N_0 = 500$ |
| Baseline area | $A_0 = L^2/N_0 = 1/500$ |
| Baseline contour length | $L_0 = \phi A_0^{1/2}$ |
| Shape index | $\phi = 3.7$<br>(hexagonal preference) |
| Elastic parameters | $K_V = 1.0 \times \eta$, $K_L = 0.1 \times \eta$ |
| Numerical time step | $\Delta t = 1.0 \times 10^{-3}$ |
| Time unit | 1 simulation time unit = 2 min |
| Simulated time | 0–500 min |

## Notes

### Competing Interest Statement

The authors have declared no competing interest.

### Summary of Updates

New Fig. 6C, S9, S12. Text revision.

